# Histone H1.0 Couples Cellular Mechanical Behaviors to Chromatin Structure

**DOI:** 10.1101/2022.11.29.518399

**Authors:** Shuaishuai Hu, Douglas J. Chapski, Natalie Gehred, Todd H. Kimball, Tatiana Gromova, Angelina Flores, Amy C. Rowat, Junjie Chen, René R. Sevag Packard, Emily Olszewski, Jennifer Davis, Christoph D. Rau, Timothy A. McKinsey, Manuel Rosa Garrido, Thomas M. Vondriska

## Abstract

Tuning of genome structure and function is accomplished by chromatin binding proteins, which determine the transcriptome and phenotype of the cell. We sought to investigate how communication between extracellular stress and chromatin structure may regulate cellular mechanical behaviors. We demonstrate that the linker histone H1.0, which compacts nucleosomes into higher order chromatin fibers, controls genome organization and cellular stress response. Histone H1.0 has privileged expression in fibroblasts across tissue types in mice and humans, and modulation of its expression is necessary and sufficient to mount a myofibroblast phenotype in these cells. Depletion of histone H1.0 prevents transforming growth factor beta (TGF-*β*)-induced fibroblast contraction, proliferation and migration in a histone H1 isoform-specific manner via inhibition of a transcriptome comprised of extracellular matrix, cytoskeletal and contractile genes. Histone H1.0 is associated with local regulation of gene expression via mechanisms involving chromatin fiber compaction and reprogramming of histone acetylation, rendering the cell stiffer in response to cytokine stimulation. Knockdown of histone H1.0 prevented locus-specific histone H3 lysine 27 acetylation by TGF-*β* and decreased levels of both HDAC1 and the chromatin reader BRD4, thereby preventing transcription of a fibrotic gene program. Transient depletion of histone H1.0 *in vivo* decompacts chromatin and prevents fibrosis in cardiac muscle, thereby linking chromatin structure with fibroblast phenotype in response to extracellular stress. Our work identifies an unexpected role of linker histones to orchestrate cellular mechanical behaviors, directly coupling cellular force generation, nuclear organization and gene transcription.

**Graphical Abstract:** 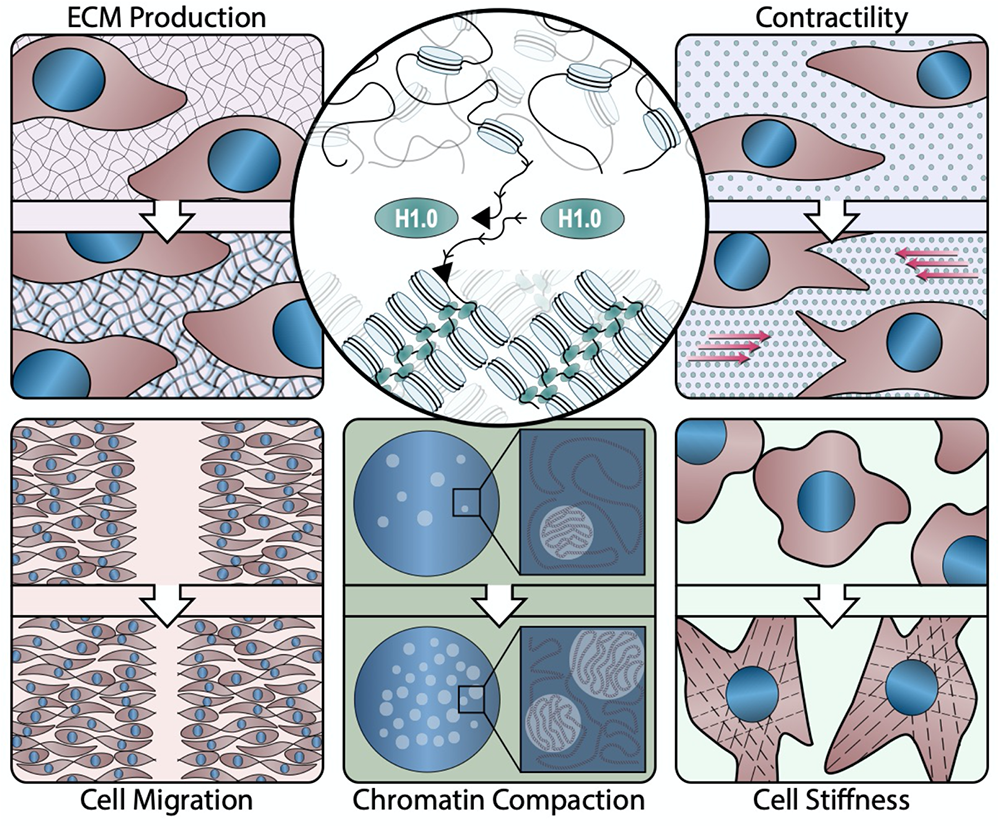

## Introduction

All metazoans have evolved processes to regulate the mechanical stability of tissues to maintain organ habitus. The varied physiological demands of different organ systems necessitate coping with a wide range of mechanical forces: teeth and bone are rigid, muscle experiences a range of tension while constantly changing its length, adipose and immune cells are respectively distensible and must deform to circulate—yet all these tissues constantly respond to external stressors. Fibroblasts are a specialized cell type present across most mammalian tissues that are responsible for synthesis of connective tissue. In adulthood, the actions of fibroblasts are essential to maintain tissue integrity and to respond to injury or cell death in various organs through a process that involves adoption of a myofibroblast phenotype. Activated myofibroblasts develop actin stress fibers, become contractile and synthesize extracellular matrix as part of a response that stiffens the tissue and heals wounds (Plikus et al., 2021). Fibroblasts are impinged upon by signals from the host tissue, facilitating transition to the activated myofibroblast phenotype. Furthermore, the plasma membrane is connected to the extracellular matrix by a network of proteins that tether the cell within the organ, thereby facilitating communication between cells and relaying extracellular physical cues to the intracellular organelles. In situations of stress, extracellular cues in the form of physical forces, cytokines and hormones induce changes in transcription that alter the mechanical properties of the cell.

The nucleus itself has been shown to directly influence distensibility of the cell and respond to mechanical signals (Uhler and Shivashankar, 2017). The nucleoskeleton is coupled to the cellular cytoskeleton, enabling force transduction and physical regulation of cellular compliance. Changes in nuclear deformability and histone post-translational modification have been shown to adaptively respond to mechanical stress, protecting the genome against aberrant gene expression (Nava et al., 2020). Changes in extracellular tension can directly impact nuclear flexibility and chromatin compaction in fibroblasts (Walker et al., 2021) and the general compaction state of chromatin can influence the mechanical stability and activation state of the cell (Kalukula et al., 2022). The functional unit of chromatin is an octamer of two copies each of histone H2A, H2B, H3 and H4 wrapped with ∼145-147 base pairs of DNA, together comprising a nucleosome (Kornberg, 1974; Luger et al., 1997). Chromatin is then packaged into higher order structures, ranging from fibers comprised of dozens of nucleosomes and a few kilobases of DNA, to topologically associated domains (thousands of nucleosomes and megabases of DNA) and nuclear territories (whole chromosomes) (Beagan and Phillips-Cremins, 2020; Dekker and Mirny, 2016; Rowley and Corces, 2018). The complex packaging rules that govern how the same genome is stored and retrieved differently across cells are known to involve the actions of histone modifying proteins, which post-translationally modify nucleosomal histones, thereby priming the targeted regions of chromatin for tasks like DNA repair, replication, transcription and gene silencing (Strahl and Allis, 2000; Wu and Grunstein, 2000). However, chromatin has unexpected functions beyond genome packaging, for example acting as a lens in rod cells of the eyes of nocturnal animals (Solovei et al., 2009) and functioning as a copper reductase (Attar et al., 2020). We reasoned chromatin may control nuclear—and thereby, cellular—compliance and set out to identify a molecular regulator of such activity. We focused on the linker histone H1 family of proteins, given their role in promoting chromatin folding (Fyodorov et al., 2018) and catalyzing formation of large spherical condensates of chromatin through phase separation (Zhang et al., 2022).

The mouse linker histone H1 family is comprised of five (H1.1-1.5) main isoforms, plus the oocyte specific H1oo, the testis specific H1t and the replacement variant H1.0. Linker histones bind the nucleosome, facilitating chromatin compaction and the formation of higher order structures comprised of multiple nucleosomes and associated DNA (Crane-Robinson, 2016). Loss of function studies have shown that individual histone H1 isoforms are dispensable for normal mouse development (Fan et al., 2001; Sirotkin et al., 1995), yet triple knockouts (deleting H1.3, H1.4 and H1.5) showed extensive developmental abnormalities (Fan et al., 2003), associated with an altered linker-core histone ratio, which was maintained when only one isoform was deleted via compensatory upregulation of other isoforms. These findings highlight the central role of histone stoichiometry in controlling normal development and tissue homeostasis (Fyodorov et al., 2018; Zhou et al., 2015). In cancer cells, altering histone H1 levels substantially reorganized global chromatin structure, decompacting topologically associating domains (Serna-Pujol et al., 2022) and shifting the genome to a more relaxed state (Yusufova et al., 2021).

Fibroblast activation leads to expression of cytoskeletal and extracellular matrix genes through a process that requires the activity of histone modifying enzymes, including histone deacetylases, and chromatin readers, including BRD4 (Felisbino and McKinsey, 2018). Furthermore, the ability of chromatin remodeling enzymes to modulate gene expression can be influenced by the local topology of chromatin (Poleshko et al., 2017), which may be regulated by the abundance of linker histone H1. A fundamental unanswered question is how the cell processes stress signals at the nucleus to remodel chromatin for precise gene expression, integrating the nucleosome-targeted actions of chromatin remodeling machinery and the genome sculping behavior of chromatin structural proteins, to elicit different mechanical responses.

We hypothesized that linker histone H1 may participate in this process of fibroblast stress response by changing the packaging of the genome. We find that levels of the histone H1 isoform, histone H1.0, are dynamic, underpinning a genome wide change in chromatin organization to facilitate transcriptional changes. We show that histone H1.0 is required for fibroblast activation in response to cytokine stimulation and overexpression of histone H1.0 is sufficient to activate fibroblasts in the absence of stimulation. Histone H1.0 acts locally to promote formation of more compact chromatin fibers and globally to condense the genome, in turn regulating cellular deformability. Histone H1.0 is required for cytokine induced reprogramming of the activating chromatin modification histone H3 lysine 27 acetylation (H3K27Ac) and acts via modulation of histone deacetylases. Histone H1.0 is also necessary for normal expression of the BET/bromodomain protein BRD4 and for its proper localization to stress responsive genes. These actions of histone H1.0 in turn regulate transcription of a cytokine-induced gene expression program involving cytoskeletal and extracellular matrix genes. Finally, we demonstrate that these chromatin regulatory actions of histone H1.0 affect a wide range of mechanical behaviors in the cell, including contractile force generation, cytoskeletal regulation, motility and extracellular matrix deposition.

## Results

### Histone H1.0 is enriched in fibroblasts and is stress responsive

To investigate the role of linker histone isoforms in response to cellular stress, we examined single cell RNA sequencing (scRNA-seq) data (Buechler et al., 2021) to understand the natural variation in these isoforms between cells and tissues. We focused on fibroblasts, ubiquitous cells present in nearly all organs of the body and involved in sensing organ level stress. Regardless of the tissue of origin, histone H1.0 is the most abundant variant in murine fibroblasts and is consistently stress activated, evidenced by abundant expression in various mouse (Figure 1A-B) and diseased human tissues (Figure 1C). Although histone H1.10 is expressed across human tissues (Figure 1C), this isoform is not conserved across mouse tissues (Figure 1A-B) and was undetectable by RT-qPCR in mouse cardiac fibroblasts. To understand how changes in extracellular tension may influence histone H1 abundance, we examined heart muscle from patients in heart failure, a condition where the muscle is under increased strain. Histone H1.0 was the primary linker variant expressed in healthy human hearts (Figure 1D) and is downregulated under conditions of increased tension in the heart (Figure 1E). Global transcriptome analyses in a genetically diverse population of mice administered the adrenergic agonist isoproterenol (Rau et al., 2015), which stiffens the muscle through increased fibrosis, showed a strong association of histone H1.0 levels with metrics of heart muscle pathology and dysfunction, including left ventricular weight (a measurement of hypertrophy) and the echocardiography parameters E and A amplitude, measurements of the heart’s ability to relax after contraction (Figure S1A). Bulk RNA-seq analyses of mouse hearts showed that despite cardiomyocytes contributing the vast majority of the heart mass, fibroblasts accounted for the greatest level of histone H1.0 transcript expression (Figure S1B). Single cell RNA-seq analyses of the mouse heart (Ren et al., 2020) showed preferential expression of the H1.0 isoform of histone H1 in fibroblasts including if all cells were considered in bulk (Figure 1F) as well as when examined across different populations of fibroblasts (Figure 1G), indicating a conserved role for the histone H1.0 isoform across most if not all cardiac fibroblasts. Based on these observations, we hypothesized that histone H1.0 may participate in fibroblast activation. To test this possibility, we adopted a primary adult mouse fibroblast model system treated with transforming growth factor beta (TGF-*β*), a cytokine involved in stress response throughout the body, including in response to tension-induced stress and fibrosis in lung (Wu et al., 2020), liver (Ding et al., 2013) and heart (Bujak and Frangogiannis, 2007). TGF-*β* treatment induced fibroblast activation as measured by periostin and alpha smooth muscle actin (*α*SMA) protein expression and demonstrated by actin stress fiber formation (Figure 1H), concomitant with a modest decrease in histone H1.0 protein levels (Figure 1I). Histone H1.2, the second most abundant isoform in fibroblasts, was increased in abundance by TGF-*β*, whereas H1.5 was decreased (Figure S1C). All other isoforms of histone H1 were unaffected by TGF-*β* treatment (Figure S1C). Thus, histone H1.0 is the primary linker histone isoform expressed in fibroblasts from various human and mouse tissues and this isoform is altered following acute and chronic stress *in vivo* and acute stress in primary cells. To understand the dynamics of the relationship between histone H1.0 levels and fibroblast phenotype, we next employed various histone H1.0 depletion strategies in primary cells.

**Figure 1.**
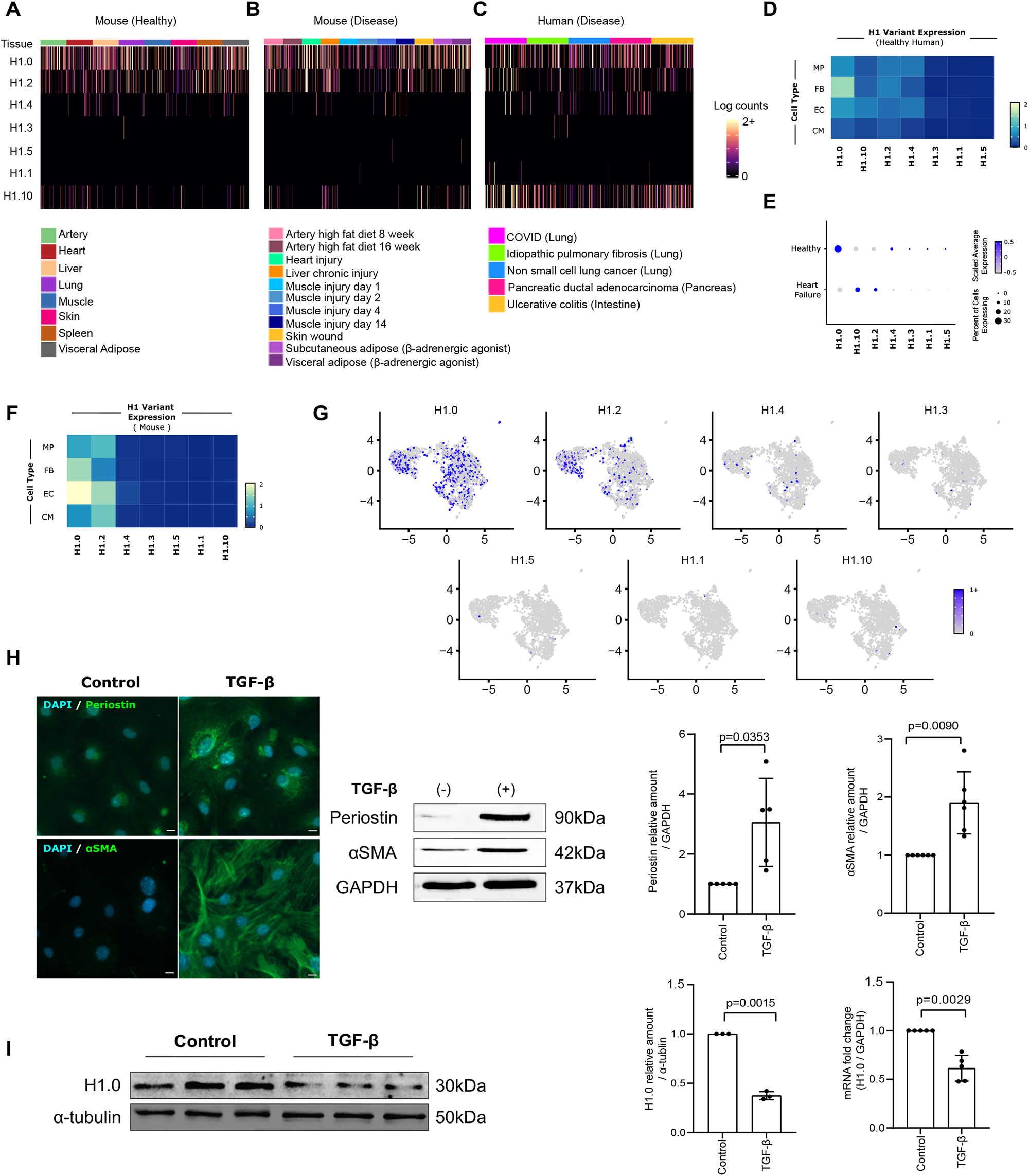
Histone H1.0 is the principal histone H1 isoform in fibroblasts. **A-C)** Heatmaps showing expression of H1 isoforms in fibroblasts from healthy murine tissue (A), murine disease models (B) and human disease tissues (C). Data from fibroXplorer.com (Buechler et al., 2021). **D)** Heatmaps showing average expression of each histone H1 isoform in macrophages (MP), fibroblasts (FB), endothelial cells (EC) and cardiomyocytes (CM) from single-cell RNA-seq analysis of healthy human hearts (Wang et al., 2020). **E)** Average expression of each H1 isoform in human cardiac fibroblasts from either healthy or heart failure samples. **F)** Heatmap of single-cell RNA-seq data (Ren et al., 2020), showing average expression of each H1 isoform in murine cardiac cell types. **G)** UMAP visualization of H1 isoforms in murine cardiac fibroblasts from single-cell RNA-seq. **H)** *Left*, Periostin and αSMA immunostaining in fibroblasts after TGF-β (10 ng/mL, 48h) or vehicle control (Nuclei stained with DAPI, scale bar=10µm; n>3). *Middle*, αSMA and Periostin expression after TGF-β treatment as measured by Western blot. *Right*, Quantification of Western blot (mean±SD; Welch’s t-test). **I)** *Left*, Western blot measurement of histone H1.0 protein abundance after fibroblast activation (72h, 10 ng/mL TGF-β). *Middle*, Immunoblot quantification of the Western blot. *Right*, RT-qPCR after TGF-β treatment (48h, 10 ng/mL; mean±SD, Welch’s t-test, n>3).

### Histone H1.0 is necessary and sufficient for fibroblast activation and controls mechanical behaviors of the cell

Previous studies demonstrated compensatory upregulation of other isoforms following germline deletion of individual linker histones (Fan et al., 2001; Sirotkin et al., 1995). We therefore used an siRNA-mediated knockdown approach targeting the six main isoforms expressed in somatic cells. Knockdown of histone H1.0 prior to administration of the cytokine was sufficient to prevent TGF-*β*-induced fibroblast activation as measured by periostin and *α*SMA transcript (Figure S1D) and protein abundance (Figure 2A). Depletion of histone H1.0 also prevented actin stress fiber formation (Figure 2B). Knockdown of histone H1.0 had minimal effects on other H1 isoforms (Figure S1F) and individual knockdown of the other five isoforms had no effect on fibroblast activation (Figure S1G-K), illustrating that even though genetic loss of linker histone H1 isoforms can be compensated developmentally (Fan et al., 2003; Fan et al., 2001), these individual isoforms play distinct roles in the somatic cell.

**Figure 2.**
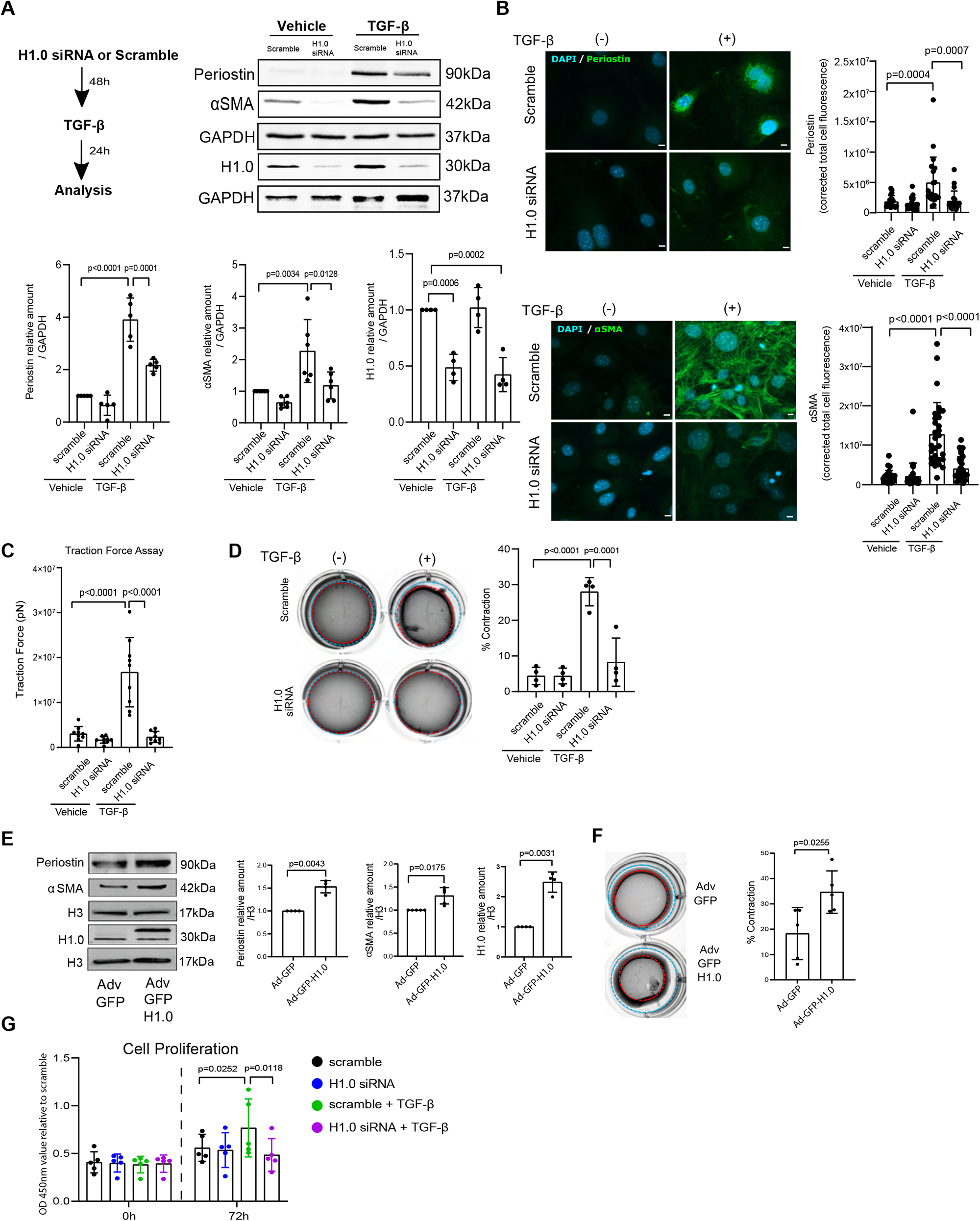
Histone H1.0 is necessary for stress-induced activation of fibroblast mechanical behaviors. **A)** Treatment schematic (*top left*). αSMA and periostin protein levels after TGF-β treatment and effect of histone H1.0 KD (*top right*; quantification, *bottom*). **B)** Immunofluorescence and quantification of periostin (*top*) and αSMA (*bottom*) protein localization *in situ* (DAPI stains cell nuclei; Scale bar=10µm. **C)** Traction force assay measuring force production at the single cell level. **D)** Gel contraction assay. **E)** *Left*, Western blot demonstrating effect of histone H1.0 overexpression on Periostin and αSMA protein levels (*right three panels*, quantitation; Welch’s t-test). **F)** Histone H1.0 overexpression and effect on gel contraction (*left*; quantification, *right*; Welch’s t-test). **G)** Absorbance measurements at 450 nm wavelength measure cell proliferation. All data are from n>3 and presented as mean ± SD with one-way ANOVA with a post-hoc Tukey test unless noted.

Depletion of histone H1.0 did not alter levels of core histones H3, H2A or H4, with only a modest change in the level of H2B (Figure S1E and S2A-C), thereby resulting in a decreased linker-core ratio. These observations are in contrast to germline knockouts of H1 isoforms, which result in compensatory alteration in other core histone levels and a maintenance of the linker to core nucleosome ratio—a key feature shown previously to regulate chromatin structure and nuclear architecture (Fyodorov et al., 2018; Zhou et al., 2015). Previous studies observed fewer linker histones per nucleosome in the setting of increased cardiac muscle tension (Franklin et al., 2012). These findings suggest that transient depletion of histone H1.0 alters the linker-core histone ratio, unmasking endogenous roles of linker histones that are compensated for in histone H1.0 germline knockouts. Interestingly, if the fibroblasts are already activated, histone H1.0 knockdown does not reverse the effects of TGF-*β* (Figure S2D) whereas simultaneous knockdown at the time of TGF-*β* treatment was sufficient to block *α*SMA but not periostin expression (Figure S2E). These observations suggest that proper histone stoichiometry is necessary for stress response in fibroblasts and that once the transcriptional program is activated in response to agonist, the window for modulating chromatin architecture to prevent this stress response has closed.

To test whether changes in marker gene expression were indicative of phenotypic changes in activated fibroblasts, we examined the effect of histone H1.0 depletion on distinct mechanical behaviors in primary cells. We employed a traction force assay, in which cells were seeded onto fluorescently labeled bovine serum albumin (BSA) beads and the deformation of the beads was used to measure the force generated by individual fibroblasts (Beussman et al., 2021). Treatment with TGF-*β* induced robust traction force generation at the individual cell level and this response was completely abrogated by depletion of histone H1.0 (Figure 2C; Note: In traction force experiments, which measure single cells, all groups were co-transfected with an siRNA conjugated to Cy3 fluorescent tag along with either the scrambled siRNA or the siRNA against histone H1.0, and only cells expressing this tag were selected for measurement). When the contractile behavior of the whole population of cells on the culture dish was examined with a gel contraction assay, histone H1.0 was again found to be necessary for the TGF-*β* induced contractile phenotype (Figure 2D). Furthermore, overexpression of histone H1.0 demonstrated that increasing the levels of this protein in the nucleus (Figure S3A) was sufficient to induce fibroblast activation in the absence of cytokine stimulation as measured by periostin and *α*SMA expression (Figure 2E) and gel contraction assay (Figure 2F). These findings demonstrate that histone H1.0 is necessary and sufficient for fibroblast activation.

To examine whether the relationship between histone H1.0 and fibroblast mechanical behaviors was more universal, we examined cells from different tissues, different species, in response to different modes of activation and by measuring distinct mechanical properties. Histone H1.0 was necessary for activation of fibroblasts from mouse lung (Figure S3B), mouse skin (Figure S3C) or human skin (Figure S3G-H). Activation of cardiac fibroblasts by angiotensin II was also dependent on histone H1.0 (Figure S3D). Depletion of histone H1.0 was sufficient to prevent cardiac fibroblast proliferation in response to TGF-*β* (Figure 2G) as measured by CCK-8 viability assay, as well as in the setting of a cell migration assay in which confluent cells are mechanically disrupted and allowed to close a pseudo wound (Figure S3E). Overexpression of histone H1.0 promoted active wound closure in the absence of TGF-*β* stimulation (Figure S3F), demonstrating that histone H1.0 is sufficient to induce this proliferative response. These data demonstrate that loss of histone H1.0 does not impair the basal ability of fibroblasts to survive or proliferate—instead, histone H1.0 is necessary and sufficient for a wide range of stress-responsive mechanical behaviors of the cell, including myofibroblast activation, proliferation, wound healing and contractile force generation. These findings indicate a conserved, tissue-independent role for histone H1.0 in regulating fibroblast stress response.

### Histone H1.0 coordinates with chromatin remodeling machinery to regulate H3K27Ac and expression of genes involved in ECM production, contraction and wound healing

To investigate molecular mechanisms whereby histone H1.0 participates in fibroblast activation, we used RNA-seq to determine the transcriptome changes following TGF-*β* treatment that are dependent on histone H1.0. Depletion of histone H1.0 prevented a select subset of transcriptional changes induced by TGF-*β* treatment (Figure 3A and Figure S4A). Ingenuity Pathway Analyses revealed that up-regulated genes whose expression was blocked by histone H1.0 knockdown are involved in key intracellular and extracellular processes (Figure 3B). KEGG analyses revealed an enrichment in pathways associated with ECM (Figure 3C and S4C) and signaling via protein kinase B/Akt (activation of which is histone H1.0-dependent (Figure S4C)), a protein associated with growth and proliferation of fibroblasts and other cells (Franke et al., 1995). Among histone H1.0 target genes was thrombospondin 4 (*Thbs4*), a secreted ECM protein known to positively regulate tissue healing and previously shown to be necessary for normal fibrotic deposition after cardiac muscle injury (Frolova et al., 2012; McLellan et al., 2020). Knockdown of histone H1.0 blocked the TGF-*β*-induced increase in THBS4 at the transcript and protein level (Figure S4D). To test whether this pathway played an essential role in fibroblast activation, we depleted *Thbs4* in cardiac fibroblasts and treated with TGF-*β*. Thbs4 was required for TGF-*β*-induced activation of myofibroblast genes periostin and *α*SMA expression (Figure S4E-F), demonstrating this to be a necessary downstream gene regulatory target of histone H1.0.

**Figure 3.**
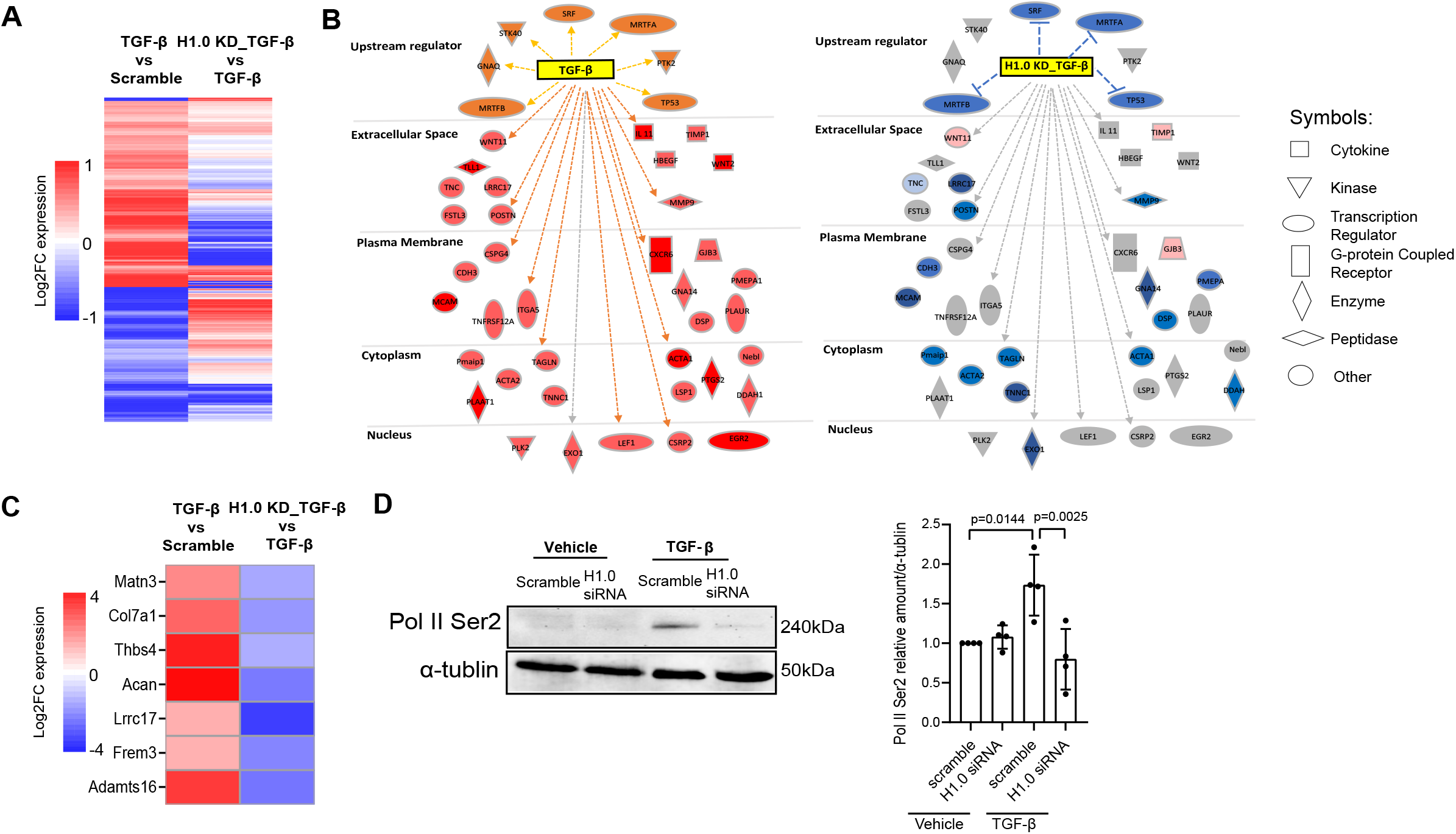
Histone H1.0 is necessary for transcriptional activation in response to TGF-β. A) Heatmap of gene expression changes TGF-β/Scramble (left) and TGF-β + H1.0 KD/TGF-β (right; p-value < 0.01). **B)** Ingenuity Pathway Analysis (IPA) identified genes significantly altered by TGF-β stimulation (*left*) and those whose expression is influenced by histone H1.0 KD prior to TGF-β (*right*). Upregulated genes are shown (*left*) (orange shading indicates activation, light/dark red indicate increased transcription following TGF-β treatment, orange lines represent predicted and measured activation). Expression of these same genes when histone H1.0 is depleted prior TGF-β versus TGF-β alone is shown on *right*, showing widespread inhibition of transcriptional changes (light/dark blue indicate decreased expression, light red indicates increased expression, and grey indicates no change). Blue lines indicate predicted and measured inhibition. Grey lines indicate no prediction of direction. **C)** Heatmap showing effect of histone H1.0 KD on the expression pattern of extracellular matrix genes activated by TGF-β treatment. **D)** Western blot shows effects of histone H1.0 KD on TGF-β-induced changes in RNA Pol II Ser2 phosphorylation. Data are mean ± SD analyzed by one-way ANOVA with a post-hoc Tukey test.

We reasoned that histone H1.0’s ability to directly bind chromatin and alter gene expression underpins these changes in gene expression and fibroblast phenotype. To test this phenomenon, we examined the influence of histone H1.0 levels on RNA polymerase II (RNAP II) expression and activation. Histone H1.0 knockdown decreased the transcript levels of RNAP II subunit a and prevented the TGF-*β*-induced increase in subunit c, whereas subunit b was slightly increased and d was unchanged by histone H1.0 depletion (Figure S4G). Histone H1.0 depletion also decreased the proportion of RNAP II that is serine 2 phosphorylated in response to TGF-*β* stimulation (Figure 3D), indicating that histone H1.0 participates in TGF-*β*-induced activation of transcription in part by regulating RNAP II subunit levels as well as post-translational modification. One mechanism of chromatin remodeling is via histone acetylation, which alters local chromatin compaction and serves to recruit reader proteins, which in turn facilitate engagement of transcriptional machinery (Shahbazian and Grunstein, 2007). Previous work has shown that inhibition of histone deacetylases (HDACs), which remove acetyl groups from lysines on histones and other proteins, is sufficient to block fibroblast activation (Travers et al., 2021) including in response to TGF-*β* (Jones et al., 2019). We thus sought to investigate whether histone H1.0 is operative in this pathway by measuring the abundance and localization of histone H3 lysine 27 acetylation (H3K27Ac), a mark associated with transcriptionally active enhancers (Creyghton et al., 2010). Depletion of histone H1.0 led to an increase in total H3K27Ac (Figure 4A), in agreement with a shift towards more active chromatin and suggesting that loss of histone H1.0 has a similar effect on the cell as inhibition of HDACs. To examine the role of histone H1.0 in regulating chromatin accessibility via acetylation, we conducted histone H3K27Ac ChIP-seq in TGF-*β*-treated cells in the presence and absence of histone H1.0. Remarkably, depletion of histone H1.0 blocks the locus specific changes in H3K27Ac induced by TGF-β (Figure 4B). Genes whose increase in H3K27Ac occupancy was blocked by histone H1.0 knockdown were enriched in pathways associated with cell migration, proliferation and ECM production (Figure 4C), demonstrating that histone H1.0 is necessary for the proper acetylation of chromatin around these stress responsive genes involved in cellular mechanical behaviors.

**Figure 4.**
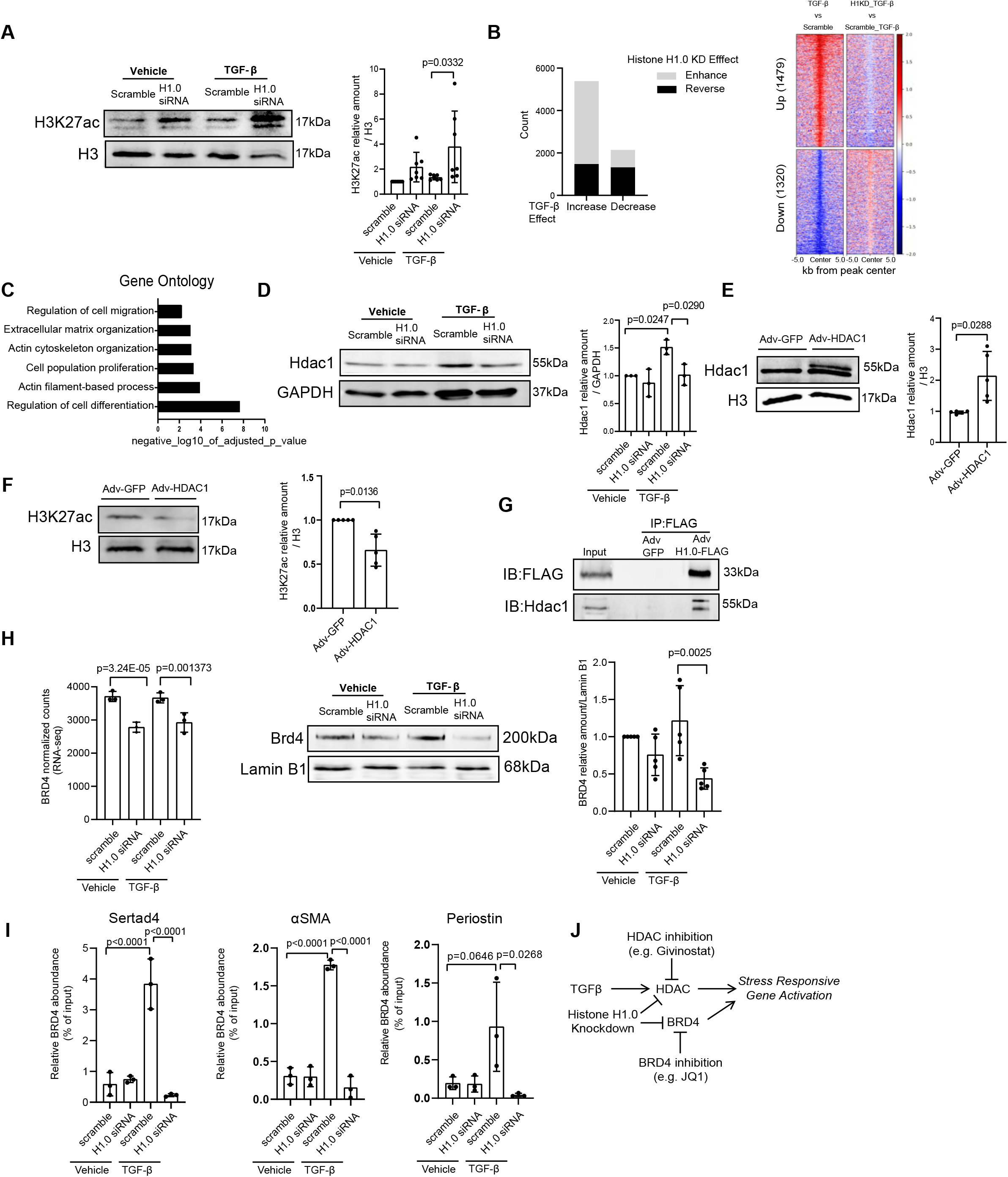
Histone H1.0 depletion prevents TGF-β-induced histone H3K27Ac and modulates actions of HDAC1 and BRD4. **A)** Immunoblot showing changes in global H3K27Ac after histone H1.0 depletion (*left*; quantification, *right*; mean ± SD, one-way ANOVA with a post-hoc Tukey test). **B)** *Left*, Stacked bar chart showing direction of histone H1.0 KD-induced H3K27ac occupancy change (H1.0 KD + TGF-β relative to TGF-β treatment alone, *y-axis*) in regions undergoing significant (FDR < 0.05) change in H3K27ac with TGF-β alone. Black and grey coloring indicate regions undergoing reversed or enhanced H3K27ac occupancy in the knockdown condition, respectively. *Right*, Heatmap depicting the log2FoldChange in H3K27ac in regions that undergo a significant increase in H3K27ac occupancy with TGF-β and are prevented with histone H1.0 KD (*top*). A visualization for the opposite behavior, regions with decreased H3K27ac occupancy that is prevented by histone H1.0 KD (*bottom*). **C)** Gene Ontology analysis of 300 unique genes closest to the regions whose increases in H3K27ac occupancy and increases in transcription after TGF-β are histone H1.0-dependent. **D)** *Left*, Western blot of HDAC1 showing effect of histone H1.0 KD on TGF-β-induced Hdac1 upregulation (r*ight*, quantification, mean ± SD, one-way ANOVA with a post-hoc Tukey test. **E)** *Left*, Fibroblasts were transfected with human Adv-HDAC1 or Adv-GFP for 48 hours and Hdac1 protein levels were detected by Western blot (*right*; quantification, mean ± SD, Welch’s t-test). **F)** *Left*, Western blot showing changes in H3K27ac abundance after Hdac1 overexpression (*right*, quantification; mean ± SD, Welch’s t-test). **G)** Co-IP assay, performed with anti-FLAG antibody using lysates from Adv-GFP or Adv-GFP-H1.0-FLAG transfected fibroblasts confirms histone H1.0 interaction with Hdac1 (representative of 5 independent Co-IP experiments). **H)** Effect of histone H1.0 knockdown on transcript (*left*, RNA-seq counts with adjusted p-values) and protein (*right*, Western blot and quantification) levels of BRD4 (mean ± SD; one-way ANOVA with a post-hoc Tukey test). **I)** ChIP-qPCR against BRD4 in primary fibroblasts examining TGF-β-induced changes in BRD4 occupancy at the promoters of *Sertad4* (*left*), αSMA (*middle*, gene name: *Acta2*), and *Periostin* (*right*) and the effects of histone H1.0 depletion. **J)** Proposed mechanism of HDAC1 and BRD4 regulation downstream of histone H1.0 signaling to control cellular stress response based on data from this paper and previous publications.

Increased pressure in the heart, which leads to fibroblast activation, pathologic muscle growth and heart failure, is associated with elevated HDAC activity (Zhang et al., 2002) and our data demonstrate that, at the molecular level, histone H1.0 is necessary for TGF-*β*-induced upregulation of HDAC1 (Figure 4D). HDAC1 is preferentially expressed in fibroblasts versus muscle cells in the heart (Nural-Guvener et al., 2014) and overexpression of HDAC1 is sufficient to diminish global levels of H3K27Ac (Figure 4E-F; HDAC inhibition has been previously shown to block TGF-*β*-induced production of ECM (Barter et al., 2010)). Co-immunoprecipitation demonstrates that histone H1.0 can bind HDAC1 in fibroblasts (Figure 4G; reciprocal immunoprecipitation yielded the same finding, *not shown*), as previously documented for other HDAC isoforms in human cell lines (Kalashnikova et al., 2013), implying this regulatory interaction can also occur at the protein level. Thus, one mechanism by which histone H1.0 can regulate transcription is by preventing upregulation of HDAC and thereby reducing its binding to, and silencing of, proliferation, migration and ECM associated genes.

To further investigate the molecular basis for how altered histone acetylation levels may influence transcription, we examined expression of the BET/Bromodomain containing chromatin reader protein BRD4, which binds acetylated lysines (Dey et al., 2003), facilitating recruitment of positive transcription elongation factor b (P-TEFb) thereby releasing transcriptional pausing (Zippo et al., 2009). Inhibition of BRD4 has been previously shown to block cell growth (Maruyama et al., 2002), fibroblast activation (Xiong et al., 2016) and neural crest development, through mechanisms that involve improper folding of chromatin (Linares-Saldana et al., 2021). We observe that histone H1.0 depletion leads to a decrease in BRD4 transcript and protein levels (Figure 4H), suggesting that reduction in the abundance of this chromatin reader—thereby preventing recruitment of transcriptional machinery to genes with acetylated histones—is part of the mechanism by which histone H1.0 inhibition prevents fibroblast activation. These observations are in agreement with histone H1.0 dependence of changes in phosphorylation of RNA Pol II at Ser2 (Figure 3D), given that BRD4 is known to recruit the essential RNA Pol II regulatory factor P-TEFb (Yang et al., 2005) and promote Ser2 phosphorylation (Devaiah et al., 2012). To further validate this mechanism, we examined binding of BRD4 to known TGF-*β* target genes using ChIP-qPCR: TGF-*β* induced robust recruitment of BRD4 to the transcription start sites of Sertad4 (Stratton et al., 2019), *α*SMA and periostin, which was completely prevented in all cases by depletion of histone H1.0 (Figure 4I). These findings, together with previous studies, indicate that histone H1.0 coordinates reorganization of H3K27Ac around TGF-*β* target genes by regulating histone acetylation via HDACs and transcription by BRD4 (Figure 4J).

### Histone H1.0 regulates global chromatin compaction and cellular deformability

Unlike transcription factors or some modified core nucleosome histones, the distribution of linker histone H1 across the genome is fairly ubiquitous (Serna-Pujol et al., 2022; Teif et al., 2020). To reveal regions of targeted regulation by histone H1.0, we performed ChIP-seq for histone H1.0 (Figure S4B) and investigated regions of relative depletion as described (Ito-Ishida et al., 2018). We find that histone H1.0 is depleted at genes undergoing altered expression following fibroblast activation, with a greater depletion observed in genes whose expression is increased (Figure 5A), suggesting that histone H1.0 eviction is associated with chromatin relaxation (Yusufova et al., 2021). We confirmed specific localization of histone H1.0 to several TGF-*β* target genes using ChIP-PCR (Figure 5B), indicating that while its genomic distribution is broad, it is not uniform. Immunoprecipitation of endogenous histone H1.0 (Figure 5A) or overexpressed, tagged histone H1.0 (Figure S4I) gave similar results (Note: We also carried out a third method of histone H1.0 ChIP-seq, using an anti-FLAG antibody to pull down the FLAG-tagged histone H1.0, which gave similar results, *not shown*). Previous studies had shown histone H1 can influence chromatin fiber condensation as measured by electron microscopy (Fan et al., 2005) and thus we directly tested the role of histone H1.0 in chromatin folding by performing nuclease digestion of genomic DNA to reveal the relative ratio of compact (nuclease inaccessible) to relaxed (nuclease accessible) DNA. In agreement with this model, overexpression of histone H1.0 increased the proportion of DNA that was compacted and thus nuclease inaccessible, whereas knockdown of histone H1.0 had the antithetical effect (Figure 5C). Interestingly, treatment with TGF-*β* shifted the genome to a more compact state and knockdown of histone H1.0 reversed this effect (Figure 5C), demonstrating that this behavior of histone H1.0 to modulate chromatin fiber accessibility is operative in the context of fibroblast activation. To test whether this is a mechanism by which histone H1.0 regulates expression of stress activated genes, we performed targeted PCR for regions of periostin and *α*SMA, demonstrating that the presence of histone H1.0 tended to compact these regions of chromatin (Figure 5D). Histone H1.0 overexpression or TGF-*β* treatment renders the periostin and *α*SMA loci less accessible (thus less DNA was recovered by PCR), whereas knockdown of histone H1.0 has the opposite effect. These findings suggest that normal levels of histone H1.0 establish a microenvironment for expression or repression of genes, such that perturbing the balance of histone H1.0 levels prevents normal stress-activated transcription.

**Figure 5.**
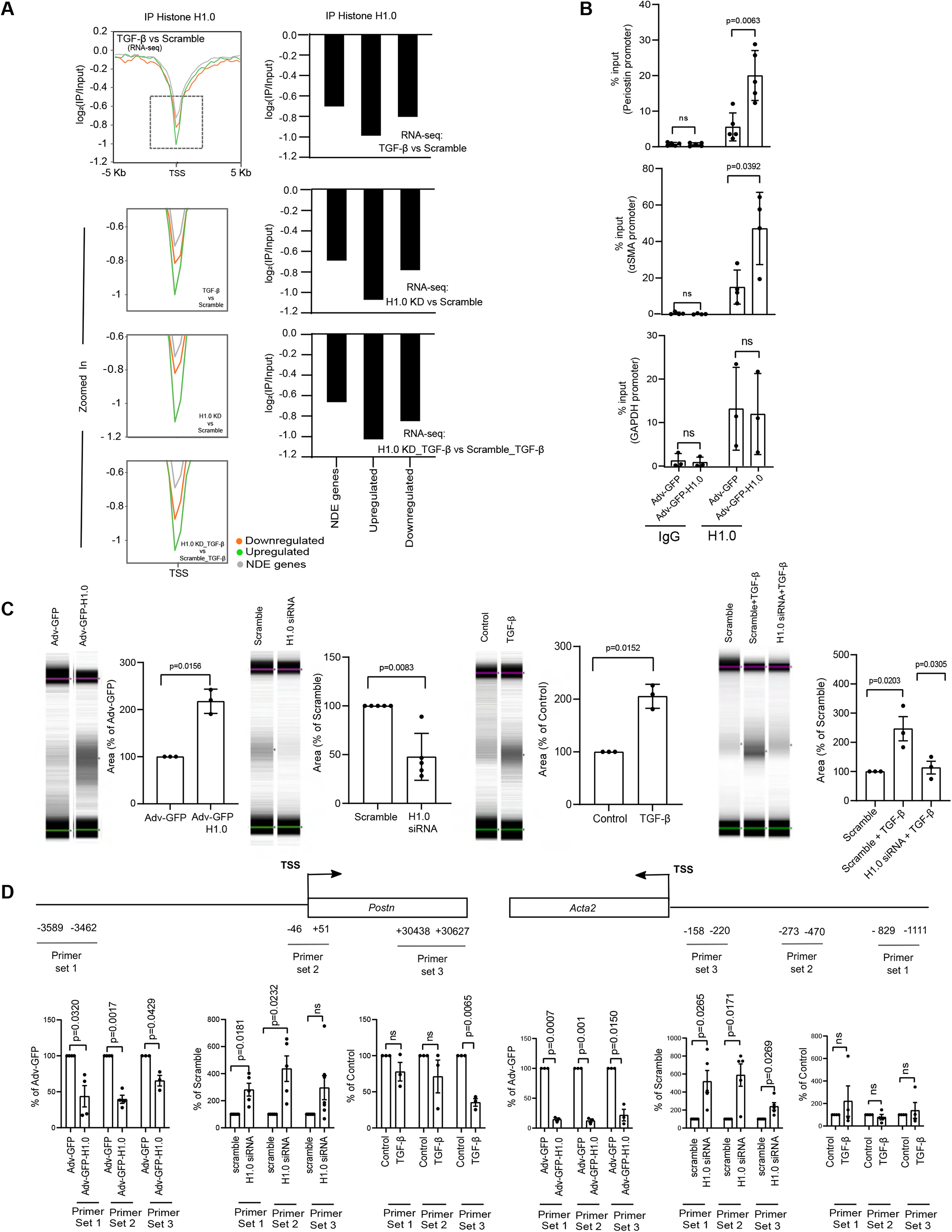
Histone H1.0 levels control chromatin fiber compaction. **A)** *Left*, ChIP-seq examining histone H1.0 occupancy at transcription start sites (TSS) relative to other genomic regions (y-axis indicates log_2_(IP/Input) signal) at differentially transcribed genes: upregulated (*green*), downregulated (*orange*) or not differentially expressed (NDE, *grey*). Labeling in bottom right of inset panel indicates the RNA-seq dataset comparisons. *Right*, Quantification of the local minimum for each condition within each inset graph from ChIP-seq profiles on *left*. **B)** ChIP-qPCR performed in isolated murine cardiac fibroblasts transfected with Adv-GFP (control) or Adv-GFP-H1.0 examining histone H1.0 occupancy in the promoter region of the *Periostin* (*top*) and αSMA (*Acta2*, *middle*; *Gapdh* promoter control, *bottom*; mean ± SD, Welch’s t-test). **C)** Gel images (upper marker 1500 bp, lower marker 25bp) of digested genomic DNA from fibroblasts transfected with Adv-GFP-H1.0 or Adv-GFP control (*first panel*), histone H1.0 siRNA or scrambled siRNA control (*second panel*), or treated with TGF-β (*third panel*). Quantification of genomic DNA in the 100-300bp range is shown next to each gel image (mean ± SD, Welch’s t-test). Effect of TGF-β in presence or absence of histone H1.0 (*fourth panel*; mean ± SD, one-way ANOVA with a post-hoc Tukey test). **D)** *Top*, location of primers used for qPCR on DNA from fibroblasts after histone H1.0 overexpression or knockdown prior to TGF-β treatment. *Bottom*, qPCR measured amount of DNA (less signal indicates less DNA and thus greater compaction of region in question; mean ± SEM, Welch’s t-test).

Nuclear deformability has been implicated in diseases such as cancer and fibrosis (Kalukula et al., 2022) and the nucleus and chromatin compaction state are major contributors to whole cell deformability (Chalut et al., 2012). We therefore sought to investigate a role for histone H1.0 to control global genome and nuclear topology, employing a cellular deformability assay (Qi et al., 2015). Depletion of histone H1.0 had a robust effect to increase the deformability of cells under basal conditions and to a lesser degree following stimulation with TGF-*β* (Figure 6A). Neither cell viability nor cell size were significantly altered by modulating histone H1.0 levels (Figure S5A-B), whereas TGF-*β* treatment increased the size and stiffness of cells independent of the nucleus (Figure S5D-F), likely contributing to the muted effect of histone H1.0 depletion on cellular deformability measurements following TGF-*β*. In contrast, increasing the abundance of nuclear histone H1.0 was sufficient to increase cellular retention in the absence of TGF-*β* (Figure 6A), which is consistent with increased cell and nuclear stiffness. Up-regulation of numerous cytoskeletal genes by TGF-*β* was blocked by depletion of histone H1.0 (Figure 6B; myosins were also under control of histone H1.0, Figure S5C) providing a mechanistic explanation for the effect of histone H1.0 depletion to alter cell compliance changes following TGF-*β*. To directly evaluate the role of histone H1.0 in genome compaction, we imaged nuclei and quantified the *chromatin condensation parameter*, a measurement of global chromatin architecture (Irianto et al., 2014), following modulation of histone H1.0 levels or hypotonic or hypertonic treatments as positive controls (which respectively decompact or compact chromatin, Figure 6C). Depletion of histone H1.0 levels caused global chromatin decondensation (Figure 6D) whereas augmentation of histone H1.0 caused condensation (Figure 6E). Taken together, these findings demonstrate that histone H1.0 modulates chromatin compaction on a genome-wide scale—directly controlling overall cell deformability—via a local mechanism in which more histone H1.0 leads to more restrictive topology (Figure 6F).

**Figure 6.**
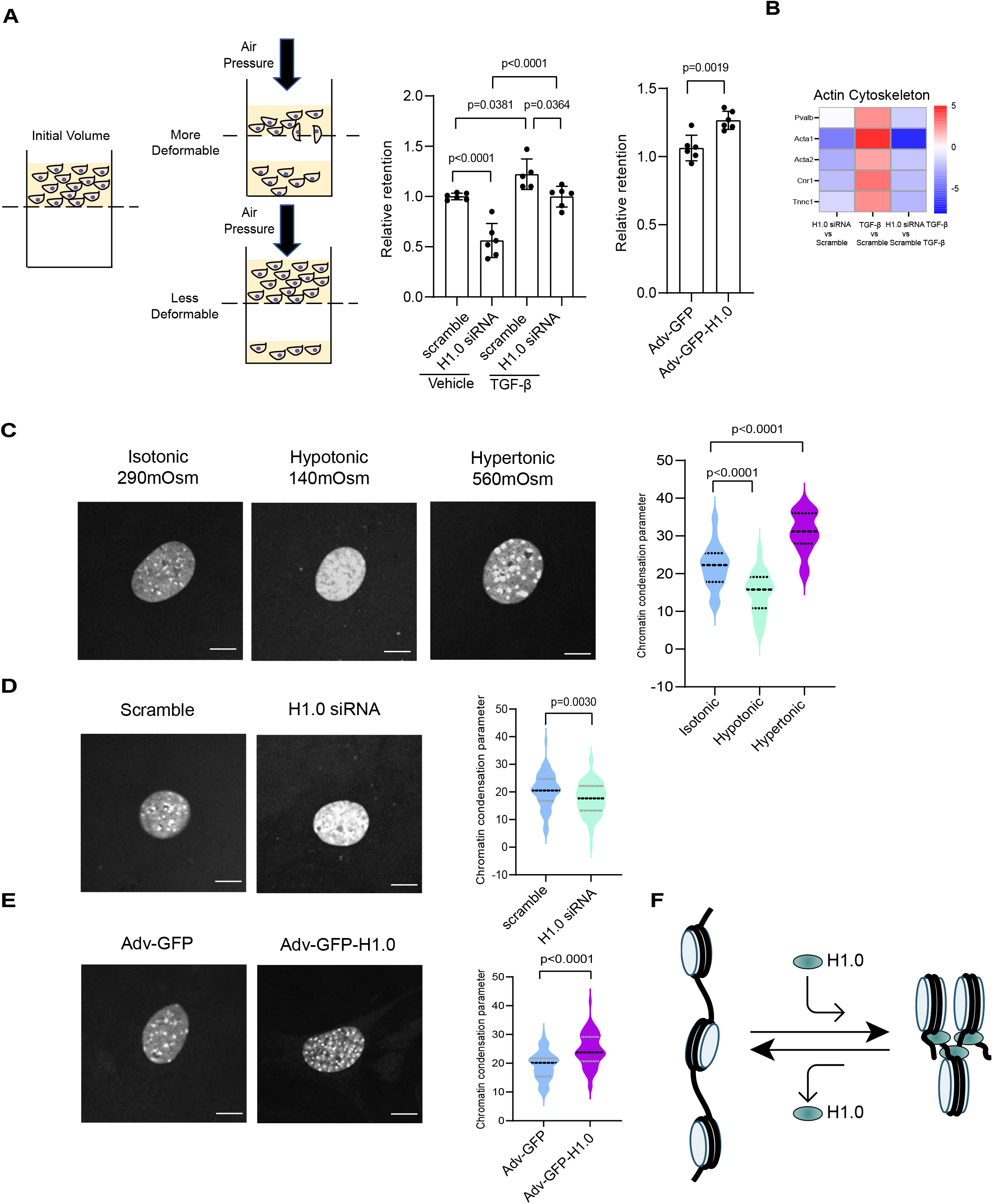
Histone H1.0 levels directly influence cellular stiffness and nuclear condensation. **A)** *Left*, Diagram of the cellular filtration assay. *Right*, Effect of histone H1.0 depletion levels and TGF-β on cellular retention (2 min applied pressure for KD and TGF-β; for H1.0 overexpressing cells, >4 min were required because of increased stiffness; mean ± SD, one-way ANOVA with a post-hoc Tukey test or Welch’s t-test). **B)** Heatmap depicting actin cytoskeleton genes and their expression after TGF-β in the presence or absence of histone H1.0. **C)** *Left*, DAPI staining of primary fibroblasts incubated under different osmotic environments for 1h (bar=10μm). *Right*, Chromatin condensation parameter quantifies condensation induced by hypotonic conditions and decondensation induced by hypertonic conditions (violin plots indicate median and quartiles; one-way ANOVA with post-hoc Tukey test, n≥20 nuclei/group). **D)** *Left*, DAPI staining of primary fibroblasts depleted of histone H1.0. *Right*, Chromatin condensation parameter, same as in C (n=70 nuclei/group). **E)** *Left*, DAPI staining of primary fibroblasts transfected with Adv-GFP-H1.0 or Adv-GFP control. *Right*, Chromatin condensation parameter, same as in C (n=70 nuclei/group). **F)** Schematic illustration of the effect of histone H1.0 abundance on chromatin compaction.

### Histone H1.0 controls fibrotic response to extracellular stress in vivo

To investigate a role for histone H1.0 to control responses to physical stress *in vivo*, we employed a model of catecholamine stimulation with isoproterenol, a nonselective *β*-adrenergic receptor agonist that increases cardiac work and thus tension on the muscle fiber, in addition to inducing fibrosis (Judd and Wexler, 1969). Notably, catecholamines can also directly increase cellular tension in non-muscle cells (Kim et al., 2019; Nguyen et al., 2016). Previous studies had shown that the fibrotic effects of isoproterenol are a complex trait strongly influenced by the genetic background of the mouse (Rau et al., 2015) and thus we chose two strains of mice to examine: C57BL/6J, which has a modest fibrotic response, and C3H/HeJ, which exhibits a more pronounced response (Figure S5G). Our analyses of C3H/HeJ mice demonstrated a positive correlation between histone H1.0 abundance and fibrotic deposition after isoproterenol treatment (Figure S5H). Administration of isoproterenol (Figure 7A) induced cardiac muscle hypertrophy in both strains, which was attenuated by coadministration of siRNA against histone H1.0 (Figure 7B). Furthermore, *in vivo* depletion of histone H1.0 blocked the isoproterenol-induced increase in the ratio of ventricular relaxation to atrial contraction (E/A) (Figure 7B), a measure of diastolic function, where a higher ratio is associated with a stiffer ventricle (Wang et al., 2019). No effect on ejection fraction, a measurement of systolic function, was observed following histone H1.0 depletion (Figure S5I). siRNA treatment was sufficient to deplete H1.0 in both mouse strains (Figure 7C & S6A) as well as to block activation of fibrotic genes, including periostin and collagen 1A1, as measured by protein/transcript abundance (Figure 7D-E). Remarkably, *in vivo* knockdown of histone H1.0 was sufficient to transiently decondense chromatin in heart muscle as measured by nuclease accessibility (Figure 7F), indicating that the *in vivo* mechanisms of protection work through actions of histone H1.0 to globally remodel the genome through local actions at the chromatin fiber. We observed a significant prevention of isoproterenol-induced fibrosis (Figure 7G), demonstrating that histone H1.0 is essential for the transcriptional program driving production of ECM *in vivo*. We also found fibrosis in the kidneys of these isoproterenol treated animals, which was partially attenuated by depletion of histone H1.0 (Figure S6B-D). Taken together, these findings demonstrate a powerful effect of histone H1.0 to regulate fibrosis *in vivo* through its actions to control chromatin packaging.

**Figure 7.**
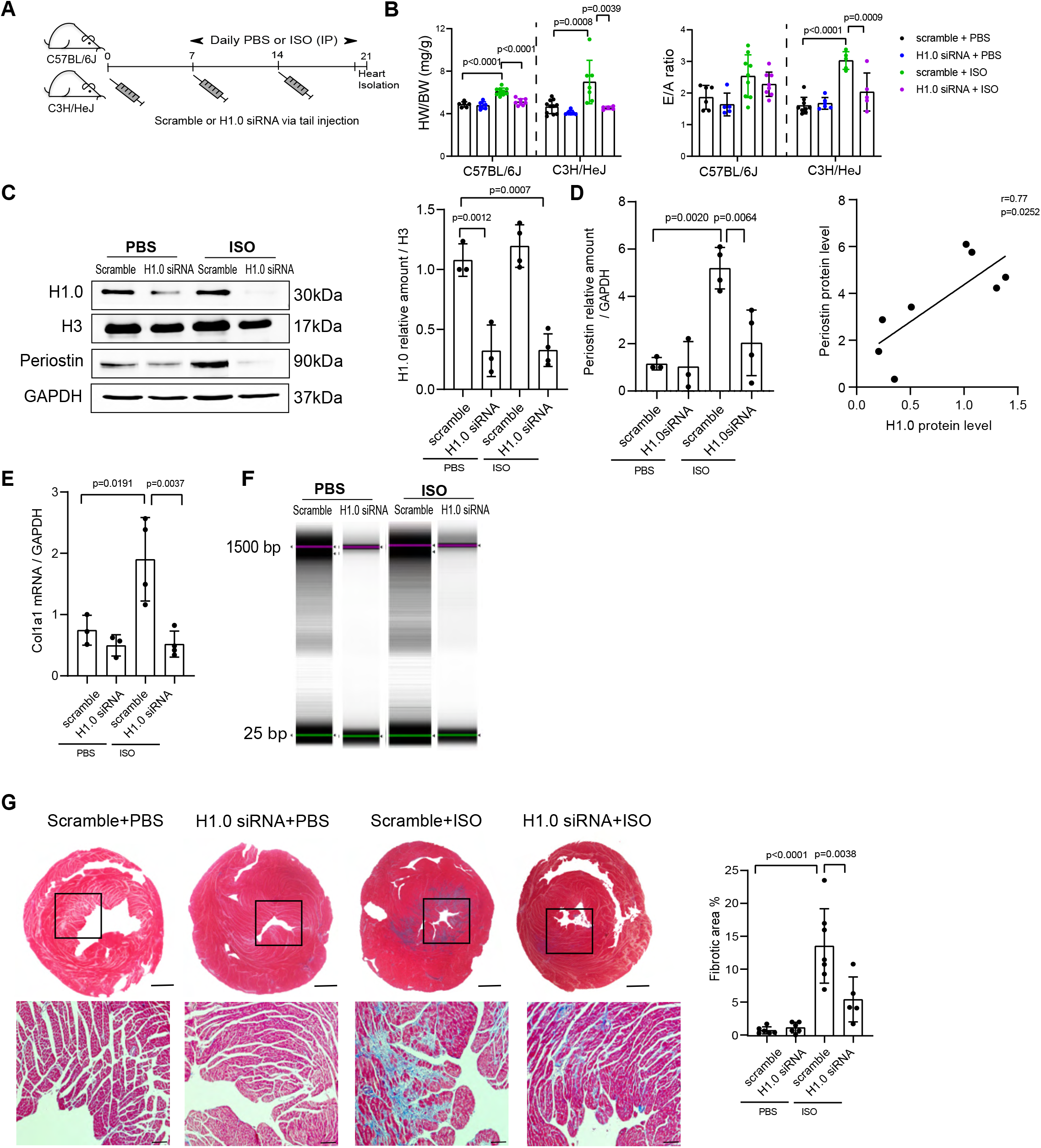
Histone H1.0 depletion prevents disease-associated cardiac fibrosis *in vivo*. **A)** Diagram of *in vivo* studies. C57BL/6J and C3H/HeJ mice were i.v. injected 3 times with siRNA targeting histone H1.0 or scrambled siRNA control on days 0, 7, and 14. At the same time, they were i.p. injected daily with isoproterenol (ISO) daily (C57BL/6J: 80mg/kg/day, C3H/HeJ: 20mg/kg/day) or PBS from day 7 to 20. **B)** *Left*, Heart weight to body weight ratios. *Right*, E/A ratios. **C)** Western blot showing histone H1.0 KD effect on ISO-induced periostin activation. **D)** Quantification of C. A positive Pearson’s correlation was observed between histone H1.0 and periostin levels in hearts from C57BL/6J mice. **E)** RT-qPCR showing effect of histone H1.0 KD on ISO-induced changes in *Col1a1* transcript abundance. **F)** Gel images of genomic DNA digested from hearts of C57BL/6J mice (n≥2/group). **G)** *Left*, Masson’s trichrome staining of heart sections to measure fibrosis (C3H/HeJ mice; bar in whole heart images = 1 mm; bar in zoomed images =100μm). *Right*, Quantification of fibrotic area. All data are presented as mean ± SD, analyzed by one-way ANOVA with post-hoc Tukey test.

## Discussion

Coupling of extracellular stress to transcription is a fundamental feature of all cells. In mammals, the various tissues of the body experience different physical forces, which require these tissues to have the ability to alter the tissue microenvironment. One manner in which this occurs is fibrosis, or the deposition of extracellular collagens and other matrix proteins which form scar to prevent tissue rupture, thereby altering parenchymal mechanics. We observe that fine tuning of histone H1 levels and chromatin compaction is necessary for response to stress stimuli and that the linker histone H1.0 isoform has a privileged role in this process. We also demonstrate gain of function experiments that show augmenting histone H1.0 levels beyond the normal stoichiometry can recapitulate the chromatin organization, gene expression and mechanical cell behaviors in the absence of changes to cellular tension or cytokine stimulation. These findings support a central role for histone H1.0 as a molecular regulator of fibroblast stress response, coupling chromatin organization with mechanical properties of the cell.

Alteration of histone H1 levels *in vivo* has been shown to shift global chromatin architecture between a relaxed (less H1) or more compact (more H1) state (Willcockson et al., 2021), and depletion of H1 (or mutations affecting the globular domain of the protein) promotes the development of lymphoma (Yusufova et al., 2021), indicating that remodeling genome packaging through histone H1 is a conserved mechanism across diseases and cell types. Our findings indicate that the levels of histone H1.0 can control chromatin condensation and thereby expression of genes associated with the cytoskeleton, force generation, ECM, and cellular motility. Concomitant with these observations, the levels of histone H1.0 control various mechanical behaviors of the cell, raising the intriguing possibility that these effects of histone H1.0 may work directly via changes in nuclear compliance—that is, to change *nuclear* stiffness as a mechanism to change *cell* stiffness—in parallel with effects of histone H1.0 to control transcription of genes associated with altering cellular rigidity. We demonstrate that the abundance of histone H1.0 on chromatin is directly associated with chromatin compaction: depleting histone H1.0 led to fiber relaxation and global decondensation, whereas overexpression had the opposite effect. These changes in chromatin accessibility prime the actions of other chromatin remodelers such as HDAC1 and BRD4, which in turn modulate transcription of stress responsive genes in a histone H1.0-dependent manner.

The enrichment of histone H1.0 we observe across fibroblast populations is in agreement with this isoform being the only polyadenylated version of the linker histone family (Doenecke et al., 1988; Kress et al., 1986) and thus, the only one likely to be strongly expressed in non-dividing cells. Deletion of histone H1.0 *in vivo* did not adversely affect mouse development (Sirotkin et al., 1995), likely due to compensation by other linker histone family members (linker to core nucleosome ratio was unchanged in these animals (Sirotkin et al., 1995)). To thoroughly explore the relationship between histone H1 isoforms and normal development and physiology, subsequent studies depleted other histone H1 isoforms by germline knockout: loss of individual isoforms H1.3, H1.4 or H1.5 failed to influence mouse development—including when combined with simultaneous loss of histone H1.0 in a double knockout model—again due to compensatory upregulation of other linker histone isoforms (Fan et al., 2001). Triple knockouts for H1.3, H1.4 and H1.5 were lethal, with no embryos surviving past E11.5 (Fan et al., 2003). While the exact stoichiometry of linker histones to nucleosome core particles at individual loci is uncertain and likely varies across the genome (and between different cell types), it has long been known that in somatic cells the average ratio approximates 1. When the linker to core ratio is maintained around 1, by altered expression of other isoforms in the setting of genetic knockout, there is no overt phenotype, whereas in the setting of triple deletions, a decreased linker to core ratio is associated with widespread developmental defects (Fan et al., 2003). This previous work indicates that linker histones compensate for each other in terms of chromatin modulation and that the cell has effective means of regulating the abundance of ancillary isoforms in the setting of deletion of individual or even multiple family members. In the present study, transient knockdown of histone H1.0 was associated with neither alterations in other linker histones nor changes in the expression of the core nucleosome histones (H2A, H3 and H4, with minimal change in H2B): thus, our intervention induced a transient decrease in the linker to core histone ratio, concomitant with perturbations in chromatin structure and the responsiveness of the cells to growth stimulus. Interestingly, extracellular stresses like TGF-*β*, or disease states involving altered muscle tension like pressure overload hypertrophy (Franklin et al., 2012), also transiently alter the abundance of linker histone H1.0 and change the linker to core ratio—suggesting that an evolutionarily conserved mechanism of stress response is a transient, reversible relaxation of chromatin modulated by linker histones.

Whether myofibroblast activation is good or bad for the organism obfuscates a more fundamental fact of this phenomenon: it is the evolutionarily conserved response to increased tissue tension and the growth factors released by this process. Our analysis of data from a recent paper using single nucleus RNA-seq to investigate dilated and arrhythmogenic cardiomyopathies (Reichart et al., 2022) revealed that histone H1.0 was the only variant increased in fibroblast subpopulations that also expressed periostin. In the same study, histone H1.0 was decreased in expression in fibroblasts from human hearts with mutation of lamin A (a nuclear membrane filament protein), which leads to cardiomyopathy. In the current investigation, histone H1.0 protein level was decreased after TGF-*β*, yet its depletion prevented fibroblast activation. It is possible that different timescales of analysis would reveal different kinetics of histone H1.0 expression: Figure 1I measures histone H1.0 expression 72h after TGF-*β* treatment, whereas the effects of knockdown are examined at 24h after TGF-*β* (72h after knockdown). Indeed, the levels of histone H1.0 vary depending on the time after TGF-*β* (*data not shown*) as well as *in vivo* following pressure overload stress to the heart (Franklin et al., 2012). Indeed, previous work has shown that levels of histone H1 variants are dynamic during development and reprogramming in the mouse embryo (Izzo et al., 2017), linking changes in global chromatin organization with the phenotypic shifts associated with maturation. Our findings suggest this dynamism of histone H1 levels may also be part of an adaptive response of fibroblasts to activation by TGF-*β* wherein the cell counteracts the activation stimulus by modulating histone levels.

Our findings support a model in which perturbation of normal chromatin architecture is an organizing feature to regulate cellular response to stress. This mechanism is centrally controlled by levels of histone H1.0 and is used by the cell to alter local chromatin compaction and to change the global stiffness of the cell to respond to altered mechanical or cytokine environment. Recent investigations have shown that linker histones participate in gene regulation through mechanisms beyond their ability to compact the chromatin fiber (Prendergast and Reinberg, 2021). The actions of histone H1.0 are not merely to turn genes on or off. We find that chromatin fibers are relaxed by histone H1.0 depletion, in agreement with previous studies of linker histones, yet histone H1.0 levels alone are not predictive of transcription. For example, the stress-activated genes periostin and αSMA, whose increase in expression is blocked by histone H1.0 depletion, are specifically bound by histone H1.0 and more compacted when it is present. Thus, during agonist stimulation, histone H1.0 dependent changes in chromatin organization must facilitate the actions of other transcriptional machinery.

We show that histone H1.0 is required for TGF-*β*-induced activation of myofibroblasts and expression of genes involved in stiffening of the extracellular matrix, a novel role for this linker histone beyond its recently identified ability to inhibit transcription of noncoding RNAs (Fernandez-Justel et al., 2022). We did not observe preferential inhibition of noncoding RNA transcription as measured by RNA-seq, potentially due to differences in measurement techniques or cell type. At the molecular level, these actions of histone H1.0 involve direct binding to chromatin and compacting local fibers to make them less accessible to transcriptional machinery. Furthermore, our data demonstrate that histone H1.0 levels influence global and gene-specific deposition of histone H3K27Ac, a mark of gene and enhancer activation. Depletion of histone H1.0 completely blocked changes in H3K27Ac induced by TGF-β (Figure 4B), including in fibrosis associated genes, notwithstanding global changes in acetylation being increased following histone H1.0 knockdown, demonstrating the specificity of this regulation. This process also involved decreased expression of HDAC1 (Figure 4D) and decreased expression of BRD4 (Figure 4H), a BET/bromodomain containing histone reader necessary for binding to acetylated histones to recruit transcriptional machinery. Small molecule based HDAC inhibition can prevent fibrosis in lung (Ren et al., 2016), liver (Liu et al., 2013), and heart (Kee et al., 2006; Lee et al., 2007; Williams et al., 2014). Similarly, small molecule based inhibition of BRD4 has independently been shown to block fibrosis across the same organs (lung (Tang et al., 2013), liver (Ding et al., 2015) and heart (Alexanian et al., 2021; Stratton et al., 2019)). In addition, previous studies in fibroblasts have shown that HDAC inhibition blocks BRD4-dependent gene activation (Travers et al., 2021). Depletion of histone H1.0 led to a decrease in HDAC levels (Figure 4D) and a global decondensation of chromatin (Figure 6D), in agreement with previous work showing that HDAC inhibition caused global chromatin decondensation and deacetylation in living cells (Strickfaden et al., 2020) and plays a critical role in genome protection during the mechanical disruptions of mitosis (Schneider et al., 2022). Histone H1 itself has been shown to be regulated by acetylation and HDAC inhibition—either by direct interaction with H1 or through chromatin decondensation—has been shown to increase histone H1 mobility on chromatin (Li et al., 2018). Both histone acetylation and BRD4 levels have been shown to drive chromatin phase separation (Gibson et al., 2019), a behavior linked to the global condensation and local fiber compaction observations made in the present study. Our findings provide a molecular link between the novel actions of histone H1.0 to regulate fibroblast mechanical behaviors and the previous observations of these histone mark erasers and readers in the setting of pathologic fibroblast activation across different organs (Figure 4J).

While histone H1.0 is dispensable for organismal development due to compensatory upregulation of other isoforms, our findings reveal a necessary role for this protein to regulate fibroblast activation and cellular stiffness in the adult mouse. Notably, knockdown of histone H1.0 alone had neither discernable effects on cell physiology in culture nor on organ function or histology *in vivo*. These observations likely represent a distinct behavior of chromatin in non-proliferating adult cells, controlled by histone H1.0: in response to stress, global chromatin changes are necessary for transcriptional activation and involve altered stoichiometry of histone H1.0, mirroring distinct effects shown for other chromatin structural proteins such as CTCF in primordial versus terminally differentiated cells (Nora et al., 2017). The actions of histone H1.0 to alter local and global compaction, H3K27Ac levels and BRD4 recruitment together facilitate priming of chromatin for transcription. These changes in chromatin architecture are responsive to stress cytokines like TGF-*β*, and hormones like angiotensin II and isoproterenol, serving to directly alter cellular stiffness by changing nuclear deformability and by ensuring proper expression of cytoskeletal, ECM and collagen genes. In this model, histone H1.0 directly links chromatin structure with cellular stress response, providing a mechanism to ensure that the microenvironment of the cell is coupled to the necessary transcriptional program to elicit distinct mechanical behaviors of the cell.

## Acknowledgements

This project was supported by National Institutes of Health (NIH) grants HL105699 (TMV) and HL150225 (TMV and TAM). TAM was also supported by NIH grants HL116848, HL147558, DK119594 and HL127240. DJC was supported by an NIH F32 (HL160099), NG by a T32 (GM007185), TG by a T32 (HL069766) and TK and MRG were supported by grants from the American Heart Association. AF and ACR were supported by grants from the National Science Foundation (BMMB-1906165, BRITE Fellow Award CMMI-2135747) and the Department of Defense (Ovarian Cancer Research Fund TEAL Expansion Award). We thank the UCLA Technology Center for Genomics & Bioinformatics for nucleotide sequencing and the Pathology Core for immunohistochemistry specimen preparation. We thank Noel Bagsik, Stacy Chang, Lesley Munoz Velazquez, Yixin Ruan and Elizabeth Soehalim for technical support in early stages of the project, Hilary Coller for human fibroblasts, Tomohiro Yokota for advice with fibrosis measurements, and Siavash Kurdistani for helpful comments on the manuscript.

## Author Contributions

SH: all experiments; AF, AR, EO, JD: cell deformability and force assays; SH, DJC, NG, TG, CDR, MRG, TMV: data analysis and interpretation; JC: *in vivo* injections; THK: microscopy, cell culture; AR, RRSP, JD, TAM, TMV: infrastructure, conceptual input and funding; SH, MRG, TMV: inception and study design; SH, DJC, NG, TMV: figures; THK: graphical abstract; TMV: writing; all authors approved the final version.

## SUPPLEMENTAL FIGURE LEGENDS

**Supplementary Figure 1. Histone H1.0 is enriched in fibroblasts, associated with cardiac dysfunction and the only isoform whose depletion influences fibroblast activation.** A) Bar chart showing significant Pearson correlation (indicated on y-axis) between cardiac H1.0 transcription in the Hybrid Mouse Diversity Panel (Rau et al., 2015) and left ventricular weight, E amplitude, and A amplitude. For all three phenotypes, p-values (top of bars) were calculated using cor.test() in R. Transcription and phenotype data were measured in hearts from control mice. B) RNA-seq data from adult mouse hearts, examining transcription in isolated myocytes, fibroblasts, and endothelial cells. C) RT-qPCR on histone H1 isoforms performed in isolated murine cardiac fibroblasts activated by TGF-β (48h, 10 ng/mL; mean ± SEM, Welch’s t-test). D) RNA-seq shows that siRNA knockdown of histone H1.0 significantly inhibits TGF-β induced αSMA (*left*) and periostin (*right*) transcription (mean±SD, adjusted p-values from RNA-seq). E) Western blotting shows histone H1.0 KD does not affect histone H3 protein level. F) Graph showing RT-qPCR of all remaining H1 isoforms after H1.0 siRNA depletion (mean±SD, Welch’s t-test for each H1 isoform). G-K) Western blots and quantitation of periostin, αSMA and the indicated histone H1 isoforms following TGF-β in the presence or absence of knockdown of the various histone H1 family members (mean ± SD, one-way ANOVA with a post-hoc Tukey test).

**Supplementary Figure 2. Effect of histone H1.0 depletion on TGF-β-dependent fibroblast activation. A-C)** Western blot and quantitation of effect of TGF-β and histone H1.0 KD on histone H2A, H2B and H4. **D)** Knockdown of histone H1.0 after fibroblast activation with TGF-β fails to block αSMA and periostin induction. **E)** Concomitant histone H1.0 KD and TGF-β treatment slightly attenuates αSMA induction. All data are mean ± SD, analyzed by one-way ANOVA with a post-hoc Tukey test.

**Supplementary Figure 3. Depletion of histone H1.0 prevents distinct fibroblast mechanical behaviors in cells from different organs and in response to different stimuli. A)** Murine cardiac fibroblasts transfected with Adv-GFP-H1.0-FLAG of Adv-GFP control (48h) were immunolabeled with an anti-FLAG antibody to show nuclear localization of histone H1.0 (*red*). DAPI (*blue*) stains cell nuclei. Scale bar=10μm (n=3 independent experiments). **B)** *Left*, Western blot in isolated murine lung fibroblasts transfected with histone H1.0 siRNA or scrambled negative control (48h) and then treated with TGF-β shows upregulation of periostin and αSMA is histone H1.0 dependent in lung. *Center left*, Immunoblot quantification. *Center right*, Efficiency of histone H1.0 KD is demonstrated by RT-qPCR (mean ± SD, one-way ANOVA with a post-hoc Tukey test). **C)** *Left*, Western blot in isolated murine skin fibroblasts transfected with histone H1.0 siRNA or scrambled negative control (48h) and then treated with TGF-β shows up-regulation of αSMA is histone H1.0 dependent in skin. *Right*, Immunoblot quantification (mean ± SD; one-way ANOVA with a post-hoc Tukey test). **D)** *Left*, Western blot in isolated murine cardiac fibroblasts transfected with histone H1.0 siRNA or scrambled negative control (48h) and then treated with Angiotensin II (48h, 1 µM) shows that induction of periostin and αSMA upregulation by Angiotensin II is histone H1.0 dependent. *Middle and right*, Immunoblot quantification (mean ± SD, one-way ANOVA with a post-hoc Tukey test). **E)** *Left*, A wound scratch assay was conducted using murine cardiac fibroblasts, demonstrating the necessary role of histone H1.0 in TGF-β-induced cell migration and proliferation. A scratch wound was made across the cell layer as indicated by the guidelines. Images were captured along the scratch wound guidelines at 0h and 24h. *Right*, Percentage of wound closure ([0h wound area – 24h wound area] / 0h wound area) was measured using ImageJ (mean ± SD, one-way ANOVA with a post-hoc Tukey test). **F)** *Left*, Wound scratch assay using murine cardiac fibroblasts transfected with Adv-GFP-H1.0-FLAG or Adv-GFP control (48h; mean ±SD, Welch’s t-test). **G)** *Left*, Histone H1.0 knockdown efficiency was confirmed by Western blot in human skin fibroblasts. *Right*, Immunoblot quantification (mean ±SD, Welch’s t-test). **H)** *Left*, Histone H1.0 knockdown impairs human skin fibroblast migration. *Right*, Quantification of the percentage of wound scratch closure ([0h wound area – 24h wound area] / 0h wound area; mean ±SD, Welch’s t-test).

**Supplementary Figure 4. Effects of histone H1.0 depletion of transcriptional machinery and stress responsive genes. A)** Principal component analysis of RNA-seq data (3 biological replicates per condition). PC1 captures effect of TGF-β, PC2 of histone H1.0 KD. **B)** Table summarizing ChIP-seq read pair counts and mapping rates for each replicate. **C)** *Left*, KEGG pathway analysis of TGF-β-upregulated genes that were histone H1.0 dependent. *Center left*, Western blot of p-Akt and total Akt in isolated murine cardiac fibroblasts following TGF-β treatment in the presence or absence of histone H1.0 KD. *Center right*, Immunoblot quantification (mean ± SD, one-way ANOVA with a post-hoc Tukey test). *Right*, Extended analysis of collagens and other ECM genes reveals dependency of their upregulation on histone H1.0. **D)** *Left*, Western blot showing that histone H1.0 KD abrogates TGF-β-induced upregulation of the extracellular matrix protein thrombospondin 4 (THBS4). *Right*, Immunoblot quantification (mean ± SD, one-way ANOVA with a post-hoc Tukey test). **E)** *Top,* Western blot confirming Thbs4 knockdown in primary cardiac fibroblasts. *Bottom*, Immunoblot quantification (mean ± SD, one-way ANOVA with a post-hoc Tukey test). **F)** *Top*, Western blot showing Thbs4 KD prevents induction of periostin and αSMA by TGF-β. *Bottom*, Immunoblot quantification (mean ± SD, one-way ANOVA with a post-hoc Tukey test). **G)** Transcript levels of four RNA Polymerase II subunits was analyzed by RNA-seq (mean ± SD, Adjusted p-values from RNA-seq). **H)** *Left*, Western blot demonstrating effect of histone H1.0 KD on TGF-β induced RNA Pol II upregulation. *Right*, Immunoblot quantification (mean ± SD, one-way ANOVA with a post-hoc Tukey test). **I)** *Left*, Histone H1.0 ChIP-seq, performed using an antibody against endogenous histone H1.0 in isolated cardiac fibroblasts overexpressing histone H1.0-FLAG, reveals depletion of H1.0 occupancy at gene TSS, with stronger depletion at upregulated genes (*green*) when compared to downregulated (*orange*) or not differentially expressed (NDE, *grey*) genes. Each inset panel label indicates the RNA-seq dataset from which each gene subset was analyzed. *Right*, Quantification of the local minimum for each condition within each inset graph from ChIP-seq profiles.

**Supplementary Figure 5. Modulation of histone H1.0 levels does not affect fibroblast viability or size. A)** Cell viability (*left*) and cell size (*right*) analyses of isolated mouse cardiac fibroblasts transfected with histone H1.0 siRNA or scrambled negative control and treated with vehicle or TGF-β show no changes among groups. **B)** Cell viability (*left*) and cell size (*right*) analyses of fibroblasts transfected with Adv-GFP-H1.0 or Adv-GFP control show no changes among groups. **C)** Heatmap depicting several myosin and focal adhesin genes whose increased expression after TGF-β is prevented by histone H1.0 knockdown. **D)** *Left*, Fibroblasts treated with vehicle or TGF-β were immunolabeled with anti-Vimentin (*green*) and anti-Lamin A/C (*red*) antibodies (*left*). DAPI (*blue*) stains cell nuclei. Scale bar=10μm. *Right*, Cell and nuclear area quantification (mean ± SD, Welch’s t-test, n≥30 nuclei from two biological replicates). **E)** *Left*, Fibroblasts transfected with histone H1.0 siRNA or scrambled negative control were immunolabeled with anti-Vimentin (*green*) and anti-Lamin A/C (*red*) antibodies. DAPI (*blue*) stains cell nuclei. Scale bar=10μm. Histone H1.0 KD promoted a reduction in nuclear area without altering cell size. *Right*, Cell and nuclear area quantification (mean ± SD, by Welch’s t-test, n≥30 nuclei from two biological replicates). **F)** *Left*, Fibroblasts transfected with Adv-GFP-H1.0 or Adv-GFP control were immunolabeled with anti-Vimentin (*red*) and Lamin A/C (*green*) antibodies. DAPI (*blue*) stains cell nuclei. Scale bar=10μm. Histone H1.0 overexpression resulted in increased cell and nuclear size. *Right*, Cell and nuclear area quantification (mean ± SD, Welch’s t-test, n≥30 nuclei from two biological replicates). **G)** Examination of fibrotic area in HMDP mouse strains undergoing isoproterenol treatment (Rau et al., 2015); asterisks indicate strains used in this study. **H)** Pearson’s correlation between histone H1.0 protein level and fibrotic area in hearts treated with ISO in the presence or absence of histone H1.0 depletion by siRNA. **H)** Neither ISO nor histone H1.0 KD affected ventricular ejection fraction in C57BL/6J and C3H/HeJ mice (mean ± SD, one-way ANOVA with a post-hoc Tukey test within strains).

**Supplementary Figure 6. Depletion of histone H1.0 prevents fibrosis *in vivo*. A)** *Left*, Efficiency of histone H1.0 KD was examined by Western blot in hearts of C3H/HeJ mice. *Right*, Immunoblot quantification (mean ± SD, one-way ANOVA with post-hoc Tukey test). **B)** *Left*, Western blot shows histone H1.0 protein levels from C57BL/6J kidney after ISO treatment and depletion of histone H1.0. *Right*, Immunoblot quantification (mean ± SD). **C)** *Left*, Representative Masson trichrome staining of C57BL/6J kidney, showing that histone H1.0 KD reduces kidney fibrosis in the ISO-treated group. Scale bars: 100 µm. *Right*, Image quantification (median and quartiles shown; one-way ANOVA with post-hoc Tukey test).

## STAR METHODS

### KEY RESOURCES TABLE

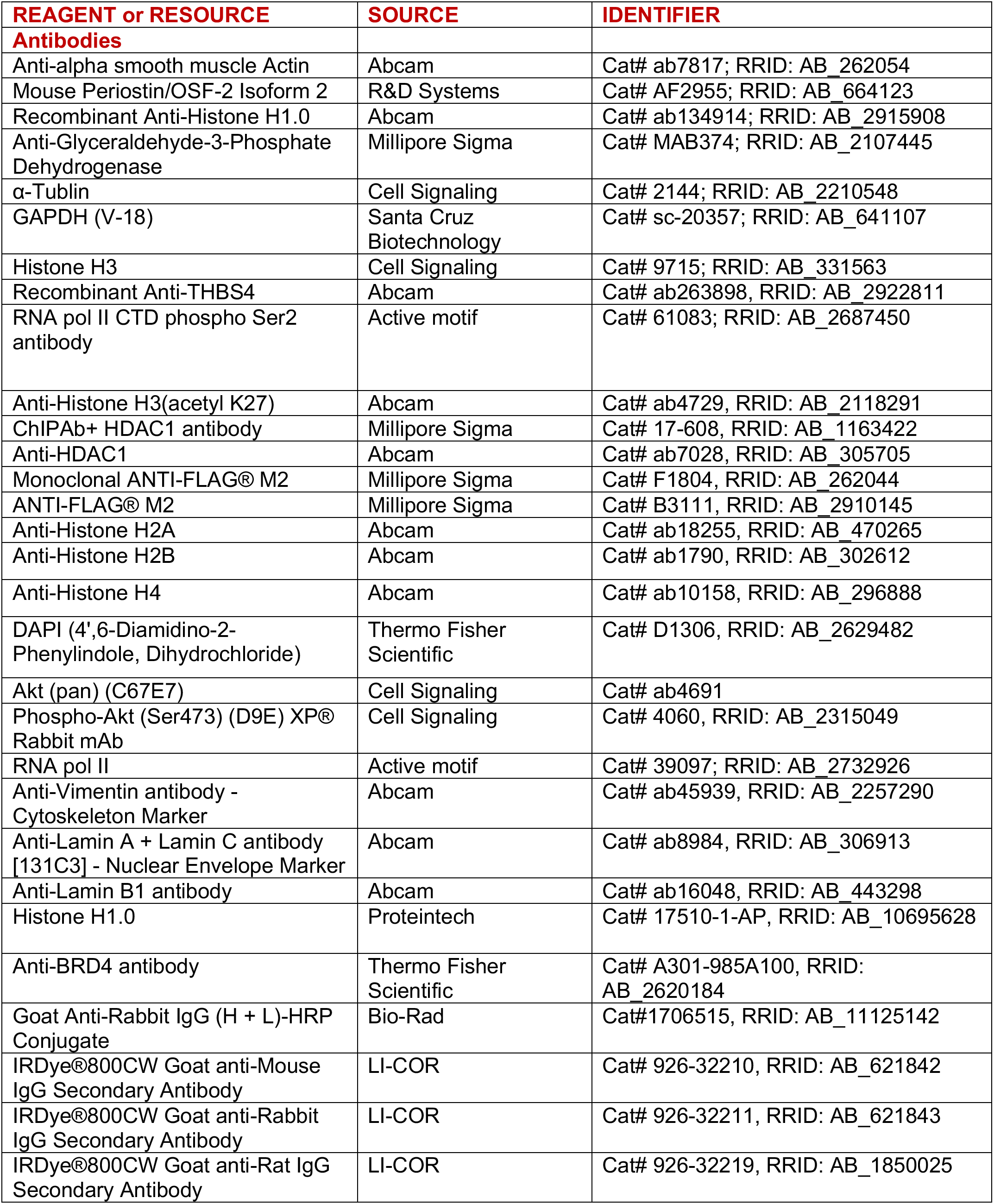
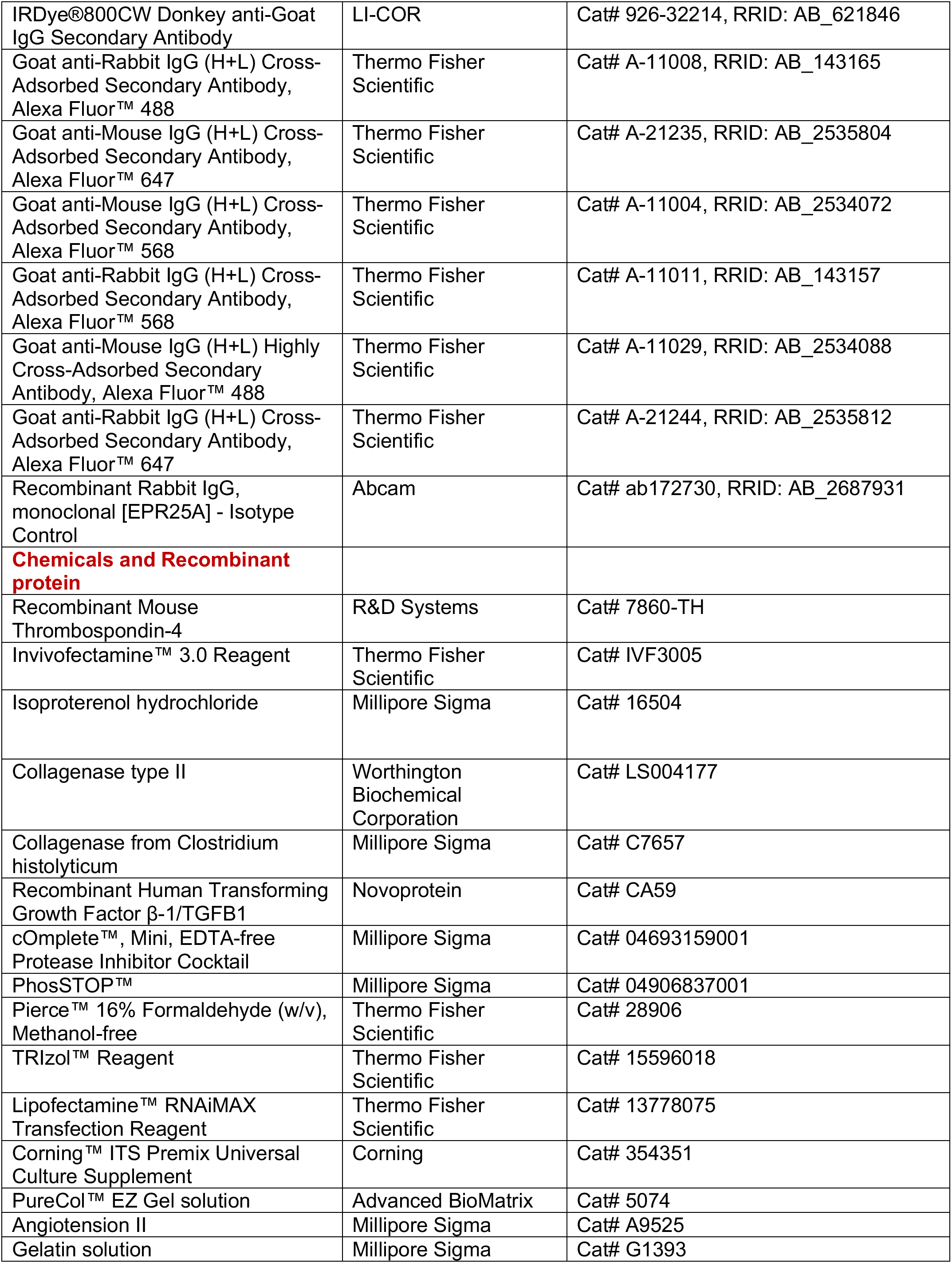
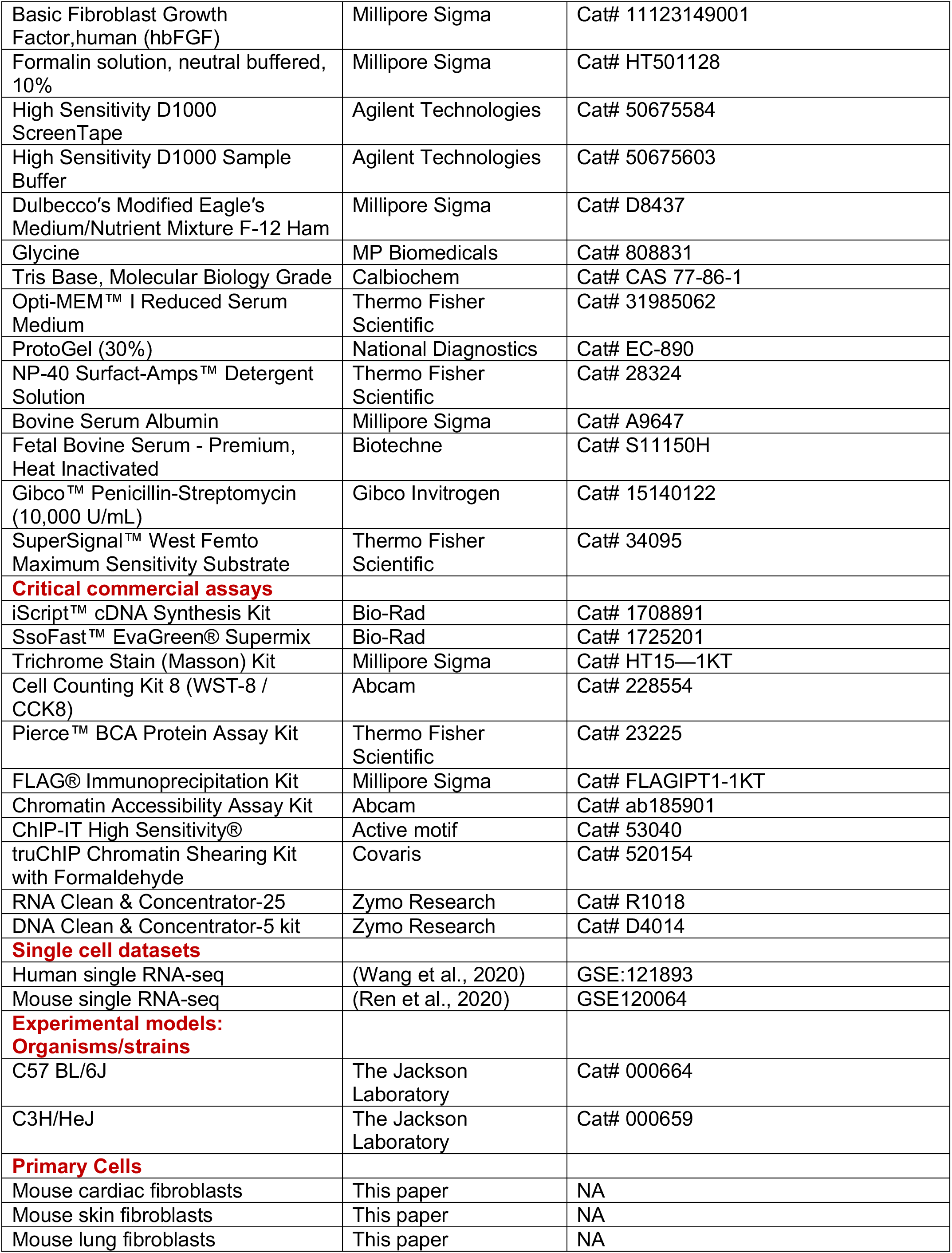
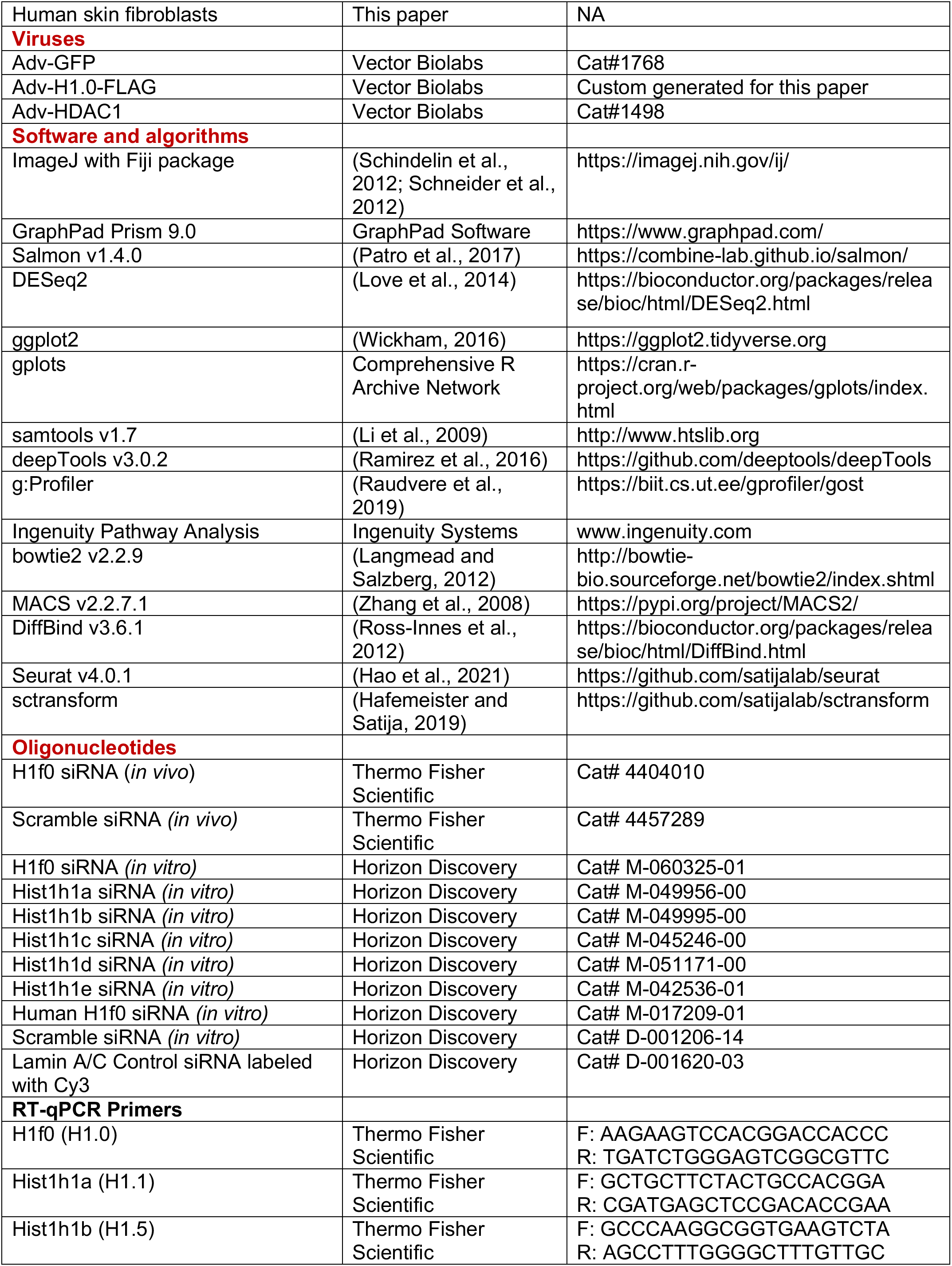
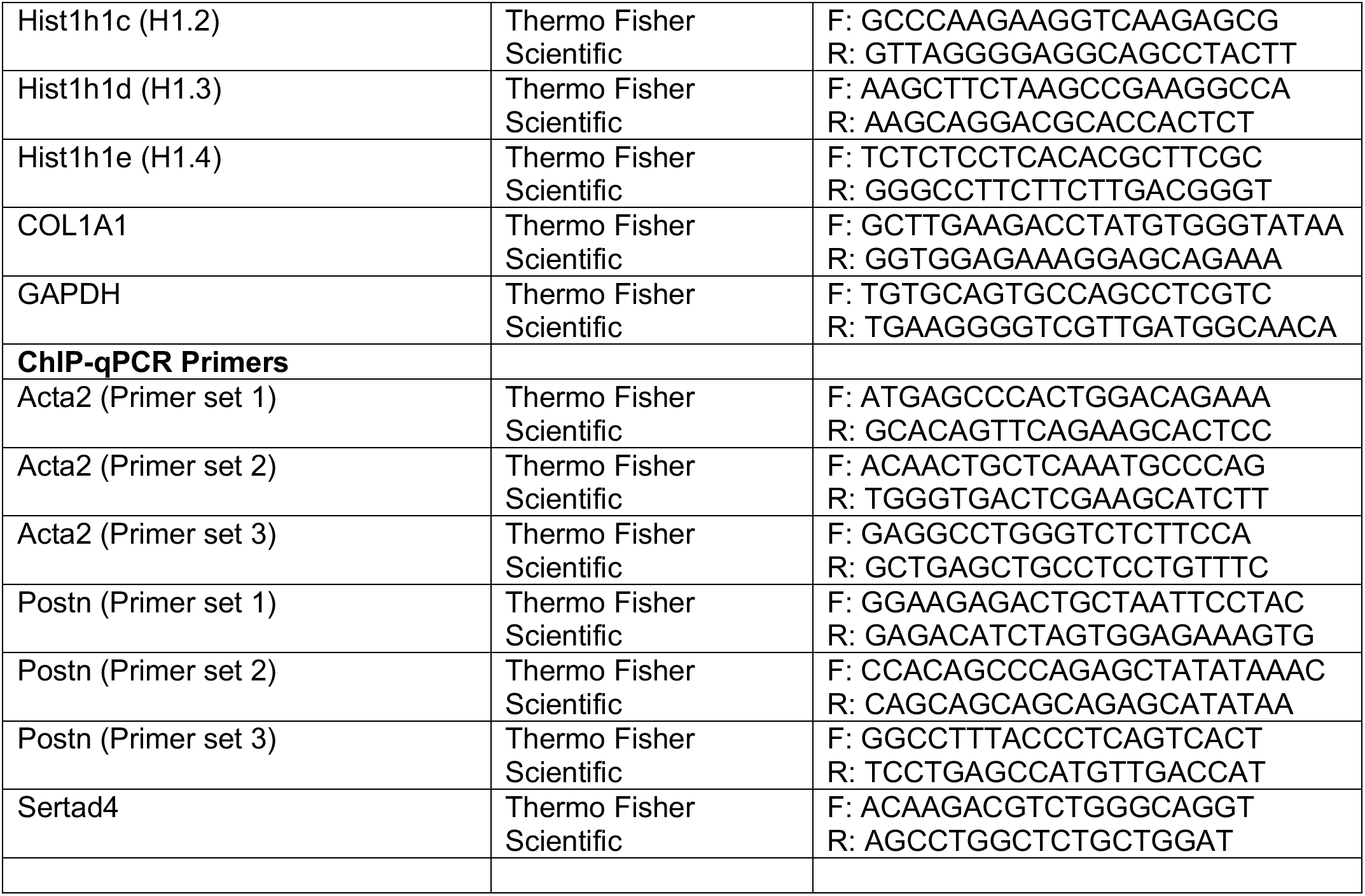

### RESOURCE AVAILABILITY

#### Lead contact

Further information and requests for resources and reagents should be directed to the Lead Contact, Thomas M. Vondriska.

#### Materials availability

All materials used in this study were obtained from commercial sources, and product details are provided in the Key Resources Table. All materials from this study are available upon reasonable request to the Lead Contact.

### EXPERIMENTAL MODELS AND SUBJECT DETAILS

#### Animal care and use

All animal studies were approved by the UCLA Animal Research Committee in compliance with the National Institutes of Health Guide for the Care and Use of Laboratory Animals. Adult female and male C57BL/6J (cat. no. 000664) and C3H/HeJ (cat. no. 000659) mice (8-12 weeks old) were obtained from Jackson Laboratory and used in the study.

#### Histone H1.0 knockdown *in vivo* in mice

H1.0 siRNA (cat. no. 4404010) and scrambled negative control siRNA (cat. no. 4457289) were purchased from Thermo Fisher Scientific. For each experimental group, siRNA (1.25 mg/kg body weight) was tail vein injected into either C57BL/6J or C3H/HeJ mice on days 0, 7, and 14 using InvivoFectamine 3.0 (cat. no. IVF3005; Thermo Fisher Scientific) according to manufacturer instructions.

#### Murine models of fibrosis

Cardiac fibrosis was induced by daily intraperitoneal injection of isoproterenol (ISO) for 14 days (C57BL/6J: 80 mg/kg/day; C3H/HeJ: 20 mg/kg/day), beginning the day of the second siRNA injection (day 7 of the *in vivo* knockdown protocol described above). Mice in negative control groups were injected daily with vehicle (phosphate-buffered saline). Mice were euthanized and then hearts, kidneys, and lungs were harvested 14 days post-ISO treatment.

#### Echocardiography

Heart function was measured by echocardiology before and after ISO treatments. Animals were anesthetized with 1.5% isoflurane and 95% O_2_ and chest hair removed. Continuous ECG monitoring was implemented, and heart rates were maintained between 400 and 500 beats per minute. Body temperature was set at 37°C using a heating pad. A Vevo 3100 imaging system was used to acquire M-mode images. LV systolic function was evaluated by calculating ejection fraction (EF%). LV diastolic function was measured by calculating the E/A ratio. All calculations were performed using the Vevo Lab 5.6.1 system.

#### Cell culture and TGF-β or Angiotensin II treatment

Adult female and male C57BL/6J primary cardiac fibroblasts were isolated using enzymatic digestion (7 mg/ml collagenase type II: cat. no. LS004177, Worthington Biochemical Corporation) followed by centrifugation (2000 rpm for 8 minutes at 4°C) and cell plating in DMEM/F12 media supplemented with 10% fetal bovine serum (FBS), 1% antibiotics (penicillin and streptomycin), and 0.1% insulin-transferrin-selenium (ITS; Corning, cat. no. 354350). After 2 hours,cells were maintained in DMEM/F12 media supplemented with 10% fetal bovine serum (FBS), 1% antibiotics (penicillin and streptomycin), human basic fibroblast growth factor (hbFGF, 1:10000 concentration from 200X stock; Millipore Sigma, cat. no. 11123149001) and 0.1% insulin-transferrin-selenium (ITS; Corning, cat. no. 354350). Media and floating cells were then removed, and fibroblasts were grown in DMEM/F12 media supplemented with 10% FBS, 1% antibiotics, hbFGF and 0.1% ITS until reaching 70-80% confluency.

Lung primary fibroblasts were isolated from adult female and male C57BL/6J mice using a previously reported method involving collagenase digestion (Edelman and Redente, 2018) and maintained in DMEM/F12 media supplemented with 20% FBS, 1% antibiotics (penicillin and streptomycin), hbFGF and 0.1% ITS until reaching 70-80% confluency.

Mouse skin fibroblasts from passage 3 to 5 and human skin fibroblasts from passage 9 were maintained in DMEM/F12 media supplemented with 10% FBS, 1% antibiotics (penicillin and streptomycin).

For all experiments involving fibroblasts, cell at 70-80% confluency was cultured in serum-free media (0.1% FBS) for 24h prior to TGF-β treatment (10 ng/mL; Novoprotein, cat. no. CA59) or Angiotensin II treatment (1µM, Sigma, cat. no. A9525).

## METHOD DETAILS

### Immunoblotting and antibodies

Protein was extracted from primary cardiac fibroblasts and lung fibroblasts using homemade RIPA lysis buffer (150mM NaCl, 5mM EDTA, 50mM Tris pH 8.0, 1% NP-40, 0.5% sodium deoxycholate, 0.1% sodium dodecyl sulfate) containing protease inhibitors (Roche, cat. no. 04693159001) and phosphatase inhibitors (Roche, cat. no. 04906837001). A homemade lysis buffer was used for protein extraction from whole heart, lung, or kidney tissue (50mM Tris pH 7.4, 10mM EDTA, 1% sodium dodecyl sulfate (SDS), 10mM sodium butyrate, 1.2mM phenylmethanesulfonyl fluoride, 1mM sodium fluoride, 1mM sodium orthovanadate) supplemented with protease inhibitor tablets (Roche, cat. no. 04693159001). Protein concentration was measured using a Pierce BCA Protein Assay (Thermo Fischer Scientific, cat. no. 23225). An equal amount of protein was loaded into an SDS-containing polyacrylamide gel. After electrophoresis, proteins were transferred to a nitrocellulose membrane (Bio-Rad, cat. no. 1620115). Membranes were blocked with 5% BSA for one hour, incubated with appropriate primary and fluorescent secondary antibodies and developed using a ChemiDoc MP Imaging System (Bio-Rad). All primary and secondary antibodies are listed in the Key Resources Table.

### Immunofluorescence

Cells were fixed at either: 1) room temperature for 10 minutes using 4% paraformaldehyde (PFA) for Figures 1 and 2; or 2) room temperature for 20 min using 1.6% PFA for Figure 6 and Supplemental Figures 5 and 6. Fixed cells were permeabilized and blocked for 1 hour using blocking buffer (5% BSA, 0.1% Triton X-100) and incubated overnight at 4°C with primary antibody: anti-αSMA (1:100, Abcam cat. no. ab7817), anti-Periostin (1:50, R&D System, cat. no. AF2966), anti-FLAG (1:100, Sigma, cat. no. B3111), anti-H1.0 (1:100, Abcam, cat. no. ab134914), anti-Vimentin (1:200, Abcam, cat. no. ab45939), or anti-Lamin A/C (1:200, Abcam, cat. no. ab8984). Appropriate concentration of secondary antibodies was incubated at room temperature for 1h. Imaging was performed at a fluorescence microscope (Zeiss Axio Vert.A1) or confocal microscope (Nikon A1R, 60x). Nuclei were stained using 4’,6-diamidino-2-phenylindole (DAPI). Secondary antibody staining alone was used as a negative control.

### Bulk RNA-seq and bioinformatics analysis

Pellets from 3 biological replicates of primary isolated cardiac fibroblasts transfected with H1.0 or scrambled siRNAs and treated with TGF-β or vehicle were sent to the UCLA Technology Center for Genomics & Bioinformatics core for RNA isolation, library preparation and sequencing. Ribosomal RNA was removed using KAPA RNA HyperPrep kit (Roche, cat. no. kk8561). Approximately 40 million paired-end reads per sample (2×150bp) were generated during sequencing. Raw fastq.gz files were downloaded and processed as described (Chapski et al., 2021), with the following modifications. Salmon v1.4.0 (Patro et al., 2017) was used to pseudoalign reads to an mm10 index built from Ensembl build 102. DESeq2 (Love et al., 2014) was used to perform differential expression testing, specifically on genes that had at least 10 reads measured between the total samples, and significantly differentially expressed genes were defined as those with adjusted pvalue (padj) less than 0.01. Principal component analysis was performed using the plotPCA() function in DESeq2 and visualized using ggplot2 (Wickham, 2016) in R. Heatmaps were visualized using gplots (https://cran.r-project.org/web/packages/gplots/index.html) in R, or Prism software v9.0 (GraphPad Software, San Diego, CA, USA). Kyoto Encyclopedia of Genes and Genomes (KEGG) pathway analysis was performed using g:Profiler (Raudvere et al., 2019) on the subset of genes upregulated by log2FoldChange of 1.5 with TGF-β (when compared to scrambled negative control) and then downregulated by log2FoldChange of 1.5 in the H1.0 siRNA + TGF-β condition (when compared to TGF-β alone).

### RT-qPCR

Total RNA was isolated from primary cardiac fibroblasts or primary lung fibroblasts from C57BL/6J mice using an RNA isolation kit (Zymo, cat. no. R1018) for RT-qPCR. Heart tissue from C57BL/6J mice was homogenized and lysed with Trizol (ThermoFischer Scientific Cat#15596018). Total RNA was extracted following with manufacturer’s instructions. cDNA was generated according to manufacturer instructions (Bio-Rad, cat. no. 1708891) and quantitative real-time PCR was performed in a CFX96 Real-Time PCR Detection System (Bio-Rad) using SsoFast EvaGreen Supermix (Bio-Rad, cat. no. 1725201). All primer sequences used in this study are listed in the Key Resources Table.

### RNAi assay *in vitro*

Lipofectamine RNAiMAX (Thermo Fisher, cat. no. 13778150) transfection was performed following the manufacturer’s protocols. Briefly, 500µL of Opti MEM Reduced Serum Media (Thermo Fischer Scientific, cat. no. 31985070) containing either Dharmacon’s siRNA targeting H1.0 (40nM, Horizon Discovery cat. no. M-060325-01), H1.1 (80nM, Horizon Discovery cat. no. M-049956-00), H1.2 (80nM, Horizon Discovery cat. no. M-045246-00), H1.3 (80nM, Horizon Discovery cat. no. M-051171-00), H1.4 (80nM, Horizon Discovery cat. no. M-042536-01), H1.5 (80nM, Horizon Discovery cat. no. M-049995-00), Thbs4 (40nM, Horizon Discovery cat. no. Cat.M-044016-01) or the appropriate concentration of siRNA negative control (Horizon Discovery cat. no. D-001206-14) were mixed with 500µL of Opti MEM Reduced Serum Media containing 20µL of Lipofectamine RNAiMAX (cat.13778075, ThermoFisher Scientific). For the human skin fibroblast experiment, transfection was performed using Dharmacon’s siRNA targeting human H1.0 (40nM, Horizon Discovery cat. no. M-017209-01). After incubating the reagents for 10 minutes at 37°C, the solution was added to the cells and slightly agitated to mix. After 24 hours incubation at 37°C, the siRNA reagent solution was removed and replaced with appropriate media according to the downstream experiment.

### Viral infection

Isolated cardiac fibroblasts were infected with either mouse Adv-GFP-H1.0-FLAG or human Adv-HDAC1, with Adv-GFP (Vector Biolabs) as a negative control, using a multiplicity of infection (MOI) of 200 plaque forming units per cell. After 24 hours incubation at 37°C, the solution was removed and replaced with DMEM/F12 media containing 10% FBS, 1% antibiotics (penicillin and streptomycin) and 0.1% ITS. For the Adv-GFP-H1.0-FLAG experiments, cells were collected 48 hours after infection. In the case of the human Adv-HDAC1 experiments, an additional infection was performed 24 hours after initial infection, and cells were collected for downstream analyses 24 hours later, for a total of 48 hours of infection.

### Collagen gel contraction assay

Primary cardiac fibroblasts were transfected with H1.0 or scrambled negative control siRNA for 48 hours. Fibroblasts suspended in 10% serum-supplemented DMEM/F-12 medium were seeded (0.5×10∧6 cells/mL) on collagen gels 24 hours prior to serum deprivation for 4 hours. At the beginning of contraction, gels were released from wells using a pipette tip and treated with TGF-β (10ng/mL; Novoprotein cat. no. CA59) for 24 hours. Primary cardiac fibroblasts transfected with Adv-GFP or Adv-GFP-H1.0-FLAG for 48 hours were suspended in 10% serum-supplemented DMEM/F-12 medium, seeded (0.5×10∧6 cells/mL) on collagen gels for 8 hours and then released from wells for 24 hours. Gel images were acquired by a ChemiDoc MP Imaging System (Bio-Rad). Gel area was calculated using ImageJ (Schneider et al., 2012) and Fiji (Schindelin et al., 2012) and contraction was reported as percentage of contraction.

### Traction force assay

Primary cardiac fibroblasts transfected with either scrambled or H1.0 siRNA with or without TGF-β stimulation were seeded onto BSA conjugated to a 647-fluorophore micropatterned onto a flexible polydimethylsiloxane (PDMS). Fibroblasts and patterned BSA dots were imaged and deformation of the dots quantified and converted into forces as described (Beussman et al., 2021).

### CCK-8 proliferation assay

Primary cardiac fibroblasts were seeded in a 48 well plate (1×10^4^ cells/well) overnight, transfected with H1.0 or scrambled siRNA for 48 hours and then treated with TGF-β (10ng/mL) or vehicle for 24 hours. After 2 hours incubation with 20µL of CCK-8 solution to each well, absorbance at 450nm wavelength was recorded in a BioTek Synergy H1 Hybrid plate reader as a readout for cell proliferation.

### Wound healing assay

Wound healing experiments were performed on 1) primary mouse cardiac fibroblasts transfected with Adv-GFP-H1.0-FLAG or Adv-GFP for 48 hours; 2) primary murine cardiac fibroblasts transfected with H1.0 or scrambled siRNA for 48 hours and then treated with TGF-β (10ng/mL) or vehicle for 24 hours; or 3) human skin fibroblasts from passage 9 transfected with H1.0 or scrambled siRNA for 48 hours. After transfection and/or TGF-β treatment, when cells were around 100% confluency, a scratch was made in the culture plate using a P200 pipette tip. Images were taken at 0 and 24 hours using a microscope (Zeiss Axio Vert.A1), and the percentage of wound closure was calculated using ImageJ (Schneider et al., 2012) and Fiji (Schindelin et al., 2012).

### Ingenuity Pathway Analysis (IPA)

Based on differential gene expression analysis of RNA-seq data detailed above, core analysis was applied in IPA to identify potential upstream regulators by comparing gene expression in Scramble +TGF-β vs Scramble and H1.0 siRNA + TGF-β vs Scramble + TGF-β groups. The activation z-score (z ≥ 2 indicates activation or z ≤ -2 indicates inhibition) was applied to predict activation or inhibition state of upstream regulators. The related gene expression changes and pathways with upstream regulators were displayed in Figure 3 to illustrate a possible mechanistic network in TGF-β treated cardiac fibroblasts. The same genes and pathways were displayed for the H1.0 siRNA + TGF-β group. The Path Designer tool within IPA was used to visualize these networks. For ease of visualization, some genes appear without lines (Ingenuity Systems).

### Immunohistology

Cardiac tissue samples were fixed in 10% formalin buffered solution (Sigma, cat. no. HT501128) overnight, dehydrated in 70% ethanol and sent to the UCLA Translational Pathology Core to generate paraffin blocks. Samples were cut into 4µm thick slices, put on slides, and stained with hematoxylin and eosin or Masson’s trichrome stain (Sigma, cat. no. HT15-1KT) to detect fibrosis. Fibrotic area for each slide was quantified and expressed as the percentage of the area occupied by the whole heart on a given slide. For kidney and lung fibrosis quantification, where the whole organ was not able to be imaged at high resolution within the same field of view, five images were taken from each mouse using 10x magnification (Zeiss Axio Vert.A1). ImageJ (Schneider et al., 2012) and Fiji (Schindelin et al., 2012) were used to calculate fibrotic area.

### Targeted nuclease digestion and RT-qPCR

Chromatin accessibility in isolated cardiac fibroblasts or whole heart from C57BL/6J mice was measured using the Chromatin Accessibility Assay Kit (Abcam, cat. no. ab185901) according to the manufacturer’s instructions. After chromatin digestion and DNA purification, a TapeStation 4200 (Agilent) was used to visualize DNA fragment size and intensity. Open chromatin is easily accessed by nucleases and digested more frequently, thereby showing a lower qPCR amplification signal relative to less accessible regions. To assess chromatin accessibility at specific loci within the periostin and αSMA promoters by RT-qPCR, three independent primer sets were designed for each promoter (sequences are provided in the Key Resources Table). Undigested DNA was used as a negative control.

### Co-immunoprecipitation (co-IP)

Primary murine cardiac fibroblasts were transfected with Adv-GFP-H1.0-FLAG or Adv-GFP for 48 hours. The co-IP assay was performed using the FLAG Immunoprecipitation kit (Sigma, cat. no. FLAGIPT1-1KT) according to the manufacturer’s instructions. Immunoprecipitated proteins were eluted using the SDS sample buffer included in the kit, and then subjected to immunoblotting.

### Hypo- and hypertonic treatment of cells

Isolated murine cardiac fibroblasts were exposed to: 1) a (1:1) mix of DMEM/F12 media supplemented with 10% FBS, 1% antibiotics (penicillin and streptomycin), and 0.1% ITS and water to reach a concentration of 140 mOsm (hypotonic treatment); or 2) a mix of DMEM/F12 media supplemented with 10% FBS, 1% antibiotics (penicillin and streptomycin), and 0.1% ITS and 10X PBS (2mL media mixed with 213μL PBS) to reach a concentration of 560 mOsm (hypertonic treatment). One hour after treatment, cells were fixed using 1.6% formaldehyde (PFA) solution in phosphate buffered saline (PBS).

### Image analysis of chromatin condensation parameter (CCP)

Primary mouse cardiac fibroblasts were fixed with 1.6% PFA and stained with DAPI. Nuclear images were taken using a confocal microscope (Nikon, A1R, 60x). Images were converted to 8-bit format and each individual nucleus was cropped from the image field by the Fiji package within ImageJ (Schindelin et al., 2012; Schneider et al., 2012). Chromatin condensation parameter (CCP) was calculated using a previously published MATLAB script (Irianto et al., 2014; Irianto et al., 2013). Briefly, the Sobel edge detection algorithm was applied to define edges within the nucleus. The density of edges within nucleus was then normalized to its cross-sectional area, giving the measured level of chromatin condensation.

### Cellular deformability assay

To measure the deformability of cardiac fibroblasts under different treatment conditions, suspended cells were filtered by air pressure through 10 µm porous membrane (Millipore) on timescales of seconds using cellular microfiltration as previously described (Qi et al., 2015). Cell deformability was determined by measuring the retention volume after 2-4 min of applied pressure. Large volume retained indicates cells are less deformable. Small volume retained indicates cells are more deformable. Prior to the filtration assay, cell viability (Trypan Blue staining) and cell size were measured by automated cell counter (TC20, Bio-Rad). To perform the assay, 400 µL cell suspension (5×10∧5/mL) were loaded into each well of a 96 well plate. The absorbance of retained cell volume was measured at 562 nm by a plate reader (SpectraMax, M2). Retention was determined by the retained volume of cells divided by the initial volume.

### ChIP-seq and bioinformatics analysis

*H1.0 and FLAG ChIP-seq*: Primary isolated murine cardiac fibroblasts were transfected with Adv-GFP-H1.0-FLAG for 48 hours. FLAG and H1.0 chromatin immunoprecipitation was performed using anti-FLAG (Sigma, cat. no. F1804) and anti-H1.0 (Proteintech, cat. no. 17510-1-AP) antibodies. In a separate experiment, primary isolated mouse cardiac fibroblasts were transfected with H1.0 siRNA or scramble for 72 hours. Chromatin shearing was performed using the truChIP Chromatin Shearing kit (Covaris, cat. no. 520154) according to the manufacturer’s instructions. DNA fragment size was assessed using a TapeStation 4200 (Agilent). Samples in the 300-500bp range were used for immunoprecipitation using the ChIP-IT High Sensitivity kit (Active Motif, cat. no. 53040). DNA was purified using a Zymo DNA Clean & Concentrator-5 kit (Zymo, cat. no. D4014). Library preparation and DNA sequencing were performed by the UCLA Technology Center for Genomics & Bioinformatics Core. Approximately 35 million paired-end reads per sample (2×150bp) were generated and used for bioinformatic analyses. Alignment of paired-end reads to the mm10 genome was performed as described (Chapski et al., 2021). After using the bamCoverage function of deepTools (Ramirez et al., 2016) with parameters --smoothLength 150 and --normalizeUsing RPGC to generate log2(IP/Input) bigWig tracks, the computeMatrix function was used with parameters -b 5000 -a 5000 --binSize 250 --nanAfterEnd --referencePoint TSS --skipZeros to calculate occupancy around transcription start sites of genes upregulated, downregulated, and unchanged with a given biological treatment. Visualization of occupancy as a ChIP-seq profile was performed using the plotProfile function of deepTools v3.0.2 with default parameters.

*H3K27ac ChIP-seq*: Three biological replicates of isolated cardiac fibroblasts from passage 1 were transfected with H1.0 or scrambled siRNAs for 48 hours and treated with TGF-β (10 ng/mL) for 24 hours.

H3K27ac immunoprecipitation was performed using the same experimental strategy as the H1.0 ChIP-seq experiment, but using an anti-H3K27ac antibody (Abcam, cat. no. ab4729). For each biological condition, a combined input sample of sonicated genomic DNA was obtained from all three biological replicates. Library preparation and paired-end sequencing were performed at the UCLA Technology Center for Genomics and Bioinformatics Core, resulting in ∼50-70 million read pairs (2×150bp) per sample. Alignment was performed against the mm10 genome using bowtie2 (Langmead and Salzberg, 2012) followed by SAM-to-BAM conversion and sorting using samtools v1.7 (Li et al., 2009).

Peak calling for each sample was performed using MACS v2.2.7.1 (Zhang et al., 2008), using the callpeak function with the following layout and parameters: --treatment ChIP_replicate.sorted.bam -- control Input_sorted.bam -f BAMPE -g mm. Differential occupancy of H3K27ac was determined using the DiffBind package (Ross-Innes et al., 2012) v3.6.1 in R, specifically on a set of consensus peaks measured in at least 3 samples across our experiment. We defined significantly differentially occupied regions as those with FDR < 0.05. To visualize H3K27ac signal in differentially occupied (FDR < 0.05) regions with TGF-β that undergo an opposite change (no thresholding) in H3K27ac signal with H1.0 depletion, we used a heatmap visualization. For each biological condition, we merged read alignments from all three biological replicates using the samtools v1.7 merge function, followed by sorting using the samtools v1.7 (Li et al., 2009) sort function and then generated genome browser tracks (bigWig files) of the log2FoldChange in signal using the bamCompare function of deepTools v3.0.2 with the -- smoothLength 150 parameter, the --outFileFormat bigwig -b1 treatment_merged.bam and -b2 control_merged.bam parameters. These bigWigs were used as inputs for the computeMatrix function of deepTools (Ramirez et al., 2016) v3.0.2 with the following parameters: reference-point --referencePoint center -b 5000 -a 5000 --skipZeros. We then used the matrix output of computeMatrix as input for the plotHeatmap function of deepTools v3.0.2 with –zMin -1.2 and --zMax 1.2, which generated the final heatmap visualization of the H3K27ac ChIP-seq data. The subset of closest genes to these regions of interest, whose expression increases (no thresholding) with TGF-β (compared to scrambled control) and decreases (no thresholding) in the H1.0 siRNA + TGF-β condition (compared to TGF-β alone) were examined by g:Profiler (Raudvere et al., 2019) using an adjusted p-value threshold of < 0.05.

### ChIP-qPCR

ChIP-qPCR against histone H1.0 or BRD4 was performed using the chromatin immunoprecipitation method described above for ChIP-seq, but with qPCR as the endpoint. For ChIP-qPCR measurements at specific loci, eluted immunoprecipitated DNA was used to perform qPCR using primer sets designed to amplify specific regions of the periostin, αSMA or Sertad4 promoters. Primer sequences and antibodies are listed in the Key Resources Table. Primers against the GAPDH promoter were used as a positive control (Active Motif, cat. no. 71018).

### Single RNA-seq bioinformatics analysis

Mouse data: Data was downloaded from NCBI (GSE120064) (Ren et al., 2020). Reads aligning to predicted genes and mitochondrial transcripts (those beginning with “Gm” and “mt-,” respectively) were removed from the UMI matrix. Seurat v4.0.1 (Hao et al., 2021) was used to create a Seurat object and perform all downstream analyses. The Seurat object was split by sample, and each of the five samples was normalized. The 3000 most variable features identified in each dataset with SCTransform (Hafemeister and Satija, 2019). The Seurat objects corresponding to each sample were then integrated together with iterative pairwise integration (Stuart et al., 2019). Briefly, SelectIntegrationFeatures was used to determine the features used to integrate the five datasets together. This function ranks features by the number of datasets in which they are deemed variable in order to prioritize features identified as highly variable in multiple datasets. This list of features was used with PrepSCTIntegration to recompute and subset the scaled feature expression data for each dataset to contain only the required anchor features for computational efficiency. Next, anchor cells (similar cells predicted to originate from a common biological state) were identified, filtered, and scored between two datasets with FindIntegrationAnchors, with dimensionality reduction by canonical correlation analysis (”cca”), normalization method set to “SCT” and the anchor features set to those identified from SelectIntegrationFeatures. Lastly, the anchors were used to integrate the datasets back together into one Seurat object with the IntegrateData function with the normalization method set to “SCT.” Default assay was changed to “integrated” before Principal Component Analysis (PCA) was performed for the first 50 principal components. An Elbow Plot was used to determine the dimensions to use (16) for identifying neighbors and clustering. The k-nearest neighbors and shared nearest neighbor graph for the dimensionality-reduced dataset was computed with FindNeighbors, with the default k.param of 20. Louvain clustering was performed with FindClusters, with a resolution of 1.2 and visualized by UMAP with UMAPPlot. Default assay was changed back to “RNA” and feature counts were normalized by cell (LogNormalize method of NormalizeData) and features centered and scaled by standard deviation (ScaleData) for downstream differential gene expression analysis. Markers of each cluster were identified with the Wilcoxon Rank Sum test via FindAllMarkers, comparing each cluster to all other cells, only testing genes that are detected in at least 25% of cells in either the cluster of interest or the other cells and only returning genes with p-value < 0.05. Cell type clusters were determined by gene expression levels of markers in Ren et al. Fig. 1F (Ren et al., 2020). The fibroblast group (”FB”, comprised of 2167 cells from clusters 5, 8, 10, and 11) was subset from the rest of the dataset. The most variable features among the fibroblasts were identified by dividing features into 20 bins based on average expression and calculating z-scores for dispersion within each bin with FindVariableFeatures(selection.method = “mvp”). PCA was performed for the first 50 principal components fibroblasts were visualized by UMAP with UMAPPlot. H1 isoform (”H1f0”, “Hist1h1a”, “Hist1h1c”, “Hist1h1d”, “Hist1h1e”, “Hist1h1b”, “H1fx”) expression levels in each cell type were compared and visualized for Figure 1 with the AverageExpression, DotPlot and FeaturePlot functions along with ggplot2. Figure 1 panel A and B heatmaps were generated from the single-cell RNA-seq data website fibroXplorer.com (Buechler et al., 2021), using 100 random cells per indicated condition.

Human data: Data was downloaded from NCBI (GSE109816 and GSE121893) (Wang et al., 2020). UMI and metadata tables from both sources were merged and intersected, respectively. Because there are cells in the UMI matrix for which metadata is not available, the UMI matrix was trimmed to include only those cells for which there is associated metadata. Per the methods (Wang et al., 2020), the UMI matrix and metadata was further filtered to include only the following: cells that express greater than 500 genes/cell, cells for which UMI count was within 2 standard deviations from the mean of log10(UMI) of all cells, cells with mitochondrial gene expression (”MT-”) less than 72%, and cardiomyocytes with sufficient UMIs (over 10000). Lastly, all mitochondrial genes (”MT-”) were filtered from the UMI matrix. Seurat v4.0.1 was used to create a Seurat object of the remaining 10,077 cells and perform all downstream analyses. The Seurat object was split by sample, and each of the 20 samples was normalized with Seurat’s NormalizeData function, in which feature counts for each cell are divided by the total counts for that cell and multiplied by 10000 before being natural-log transformed using log1p. The most variable features in each dataset were identified by dividing features into 20 bins based on average expression and calculating z-scores for dispersion within each bin with FindVariableFeatures (selection.method = “mvp”). These variable features were then used for iterative pairwise integration (Stuart et al., 2019). Briefly, SelectIntegrationFeatures was used to determine the features used to integrate the 20 datasets together. This function ranks features by the number of datasets in which they are deemed variable to prioritize features identified as highly variable in multiple datasets. Next, anchor cells (similar cells predicted to originate from a common biological state) were identified, filtered, and scored between two datasets with FindIntegrationAnchors, with dimensionality reduction by canonical correlation analysis (”cca”), normalization method set to “LogNormalize,” and the anchor features set to those identified from SelectIntegrationFeatures. Lastly, the anchors were used to integrate the datasets back together into one Seurat object with the IntegrateData function. Default assay was changed to “integrated” before PCA was performed for the first 50 principal components. The k-nearest neighbors and shared nearest neighbor graph for the dimensionality-reduced dataset was computed with FindNeighbors, with 10 dimensions used and the default k.param of 20. Louvain clustering was performed with FindClusters, with a resolution of 1 and visualized by UMAPPlot. To identify fibroblast cells, the default assay was changed back to “RNA” and feature counts were normalized by cell (LogNormalize method of NormalizeData) and features centered and scaled by standard deviation (ScaleData) for downstream differential gene expression analysis. Differential gene expression in each cluster was identified with the Wilcoxon Rank Sum test via FindAllMarkers, comparing each cluster to all other cells, only testing genes that are detected in at least 20% of cells in either the cluster of interest or the other cells, and only returning genes with p-value < 0.05. Cell type clusters were determined by gene expression levels of markers in Wang et al. Fig. 1D (Wang et al., 2020). H1 isoform (”H1F0”, “HIST1H1A”, “HIST1H1C”, “HIST1H1D”, “HIST1H1E”, “HIST1H1B”, “H1FX”) expression levels in each cell type were compared and visualized for Figure 1 with the AverageExpression and DotPlot functions along with ggplot2. The fibroblast group (”FB”, comprised of 932 cells from clusters 7 and 9) were subset from the rest of the dataset for Figure 1E generation. The Figure 1 panel C heatmap was generated from the single-cell RNA-seq data website fibroXplorer.com (Buechler et al., 2021), using 100 random cells per indicated condition.

### Quantification and statistical analysis

Data are presented as the mean ± SD, unless otherwise indicated in the figure legends. Statistical analyses were performed using Prism software v9.0 (GraphPad Software, San Diego, CA, USA) using Welch’s t-test between two groups and one-way ANOVA with Tukey’s multiple comparison analysis between three or more groups. A p-value less than 0.05 was considered statistically significant.

