## Supplemental Figures for "Histone H1.0 Couples Cellular Mechanical Behaviors to Chromatin Structure"

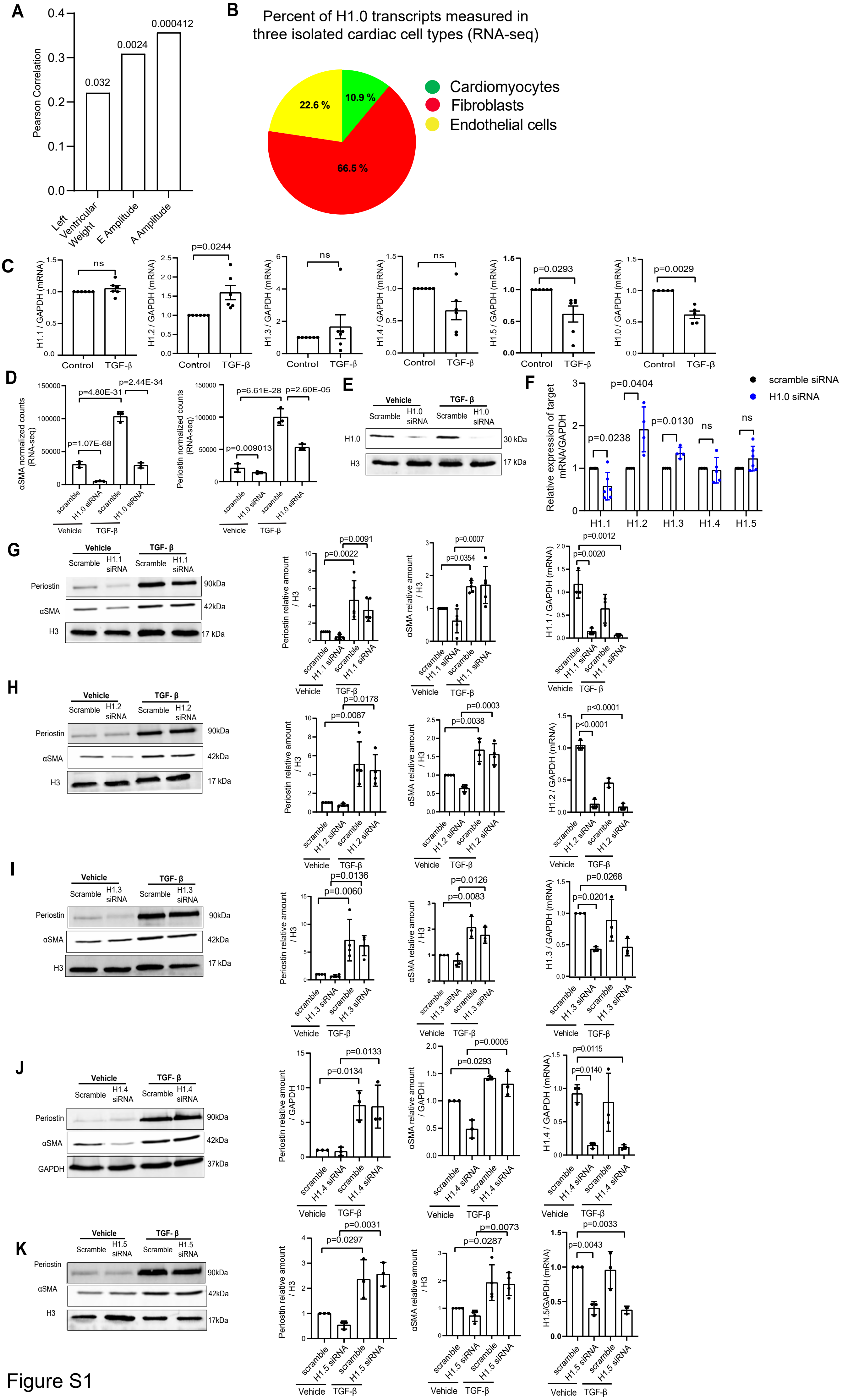

A
Pearson Correlation
0.032
0.0024
0.000412
Left Ventricular Weight
E Amplitude
A Amplitude
B
Percent of H1.0 transcripts measured in three isolated cardiac cell types (RNA-seq)
10.9 %
22.6 %
66.5 %
Cardiomyocytes
Fibroblasts
Endothelial cells
C
H1.1 / GAPDH (mRNA)
H1.2 / GAPDH (mRNA)
H1.3 / GAPDH (mRNA)
H1.4 / GAPDH (mRNA)
H1.5 / GAPDH (mRNA)
H1.0 / GAPDH (mRNA)
Control
TGF-β
ns
p=0.0244
p=0.0293
p=0.0029
D
αSMA normalized counts (RNA-seq)
Periostin normalized counts (RNA-seq)
p=1.07E-68
p=4.80E-31
p=2.44E-34
p=0.009013
p=6.61E-28
p=2.60E-05
scramble
H1.0 siRNA
Vehicle
E
Vehicle
TGF- β
Scramble
H1.0 siRNA
H1.0
H3
30 kDa
17 kDa
F
Relative expression of target mRNA/GAPDH
scramble siRNA
H1.0 siRNA
p=0.0238
p=0.0404
p=0.0130
ns
H1.1
H1.2
H1.3
H1.4
H1.5
G
Periostin
αSMA
H3
90kDa
42kDa
17 kDa
H1.1 siRNA
Periostin relative amount / H3
αSMA relative amount / H3
p=0.0022
p=0.0091
p=0.0354
p=0.0007
p=0.0020
p=0.0012
H
H1.2 siRNA
p=0.0087
p=0.0178
p=0.0038
p=0.0003
p<0.0001
I
H1.3 siRNA
p=0.0060
p=0.0136
p=0.0083
p=0.0126
p=0.0201
p=0.0268
J
H1.4 siRNA
GAPDH
37kDa
Periostin relative amount / GAPDH
αSMA relative amount / GAPDH
p=0.0134
p=0.0133
p=0.0293
p=0.0005
p=0.0140
p=0.0115
K
H1.5 siRNA
17kDa
H1.5/GAPDH (mRNA)
p=0.0297
p=0.0031
p=0.0287
p=0.0073
p=0.0043
p=0.0033


Figure S1

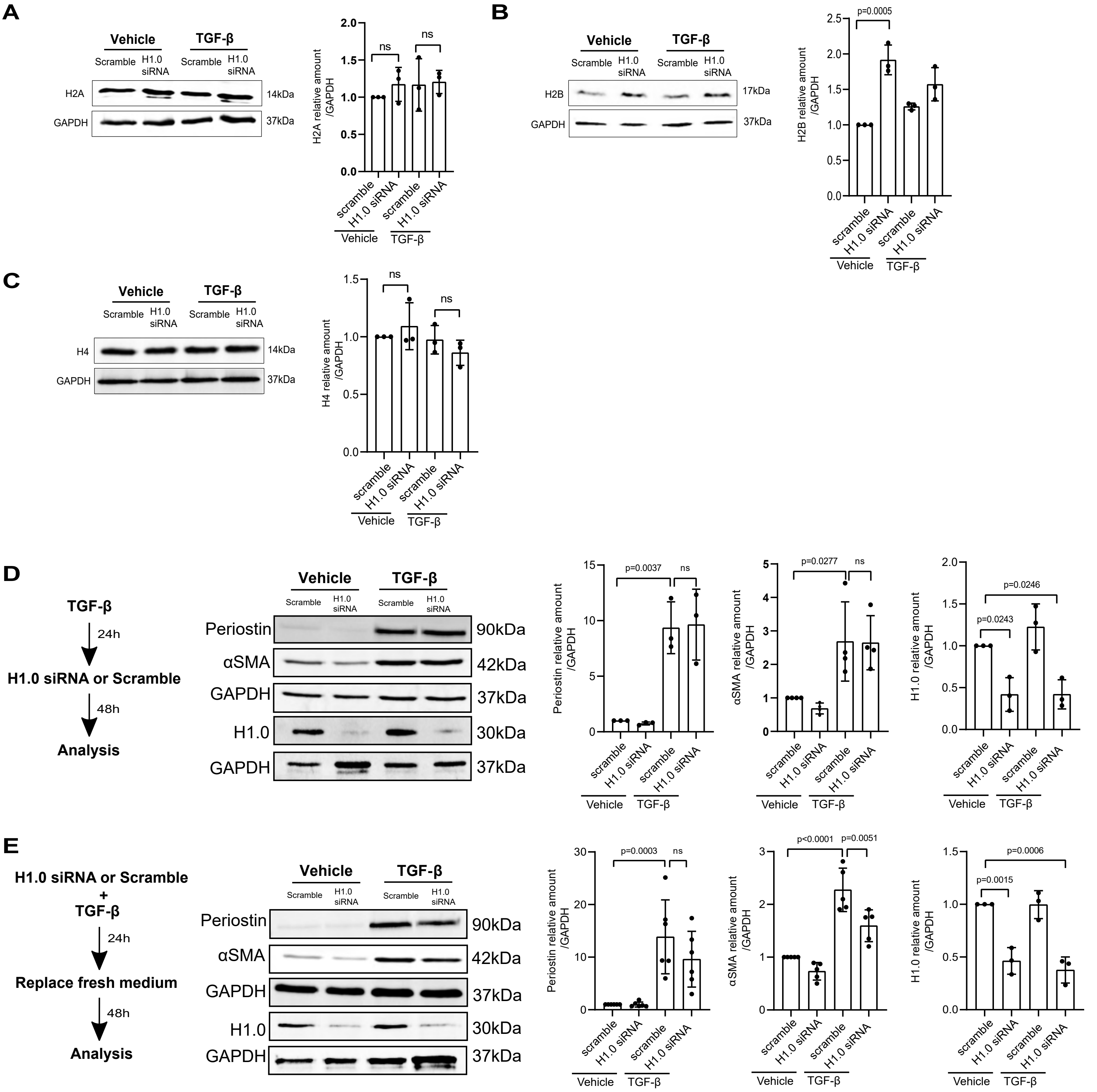
A
Vehicle
TGF-β
Scramble
H1.0 siRNA
H2A
14kDa
GAPDH
37kDa
H2A relative amount /GAPDH
ns
B
p=0.0005
H2B
17kDa
H2B relative amount /GAPDH
C
H4
H4 relative amount /GAPDH
D
TGF-β
24h
H1.0 siRNA or Scramble
48h
Analysis
Periostin
90kDa
αSMA
42kDa
H1.0
30kDa
p=0.0037
p=0.0277
p=0.0246
p=0.0243
Periostin relative amount /GAPDH
αSMA relative amount /GAPDH
H1.0 relative amount /GAPDH
scramble
E
H1.0 siRNA or Scramble + TGF-β
Replace fresh medium
p=0.0003
p<0.0001
p=0.0051
p=0.0006
p=0.0015

Figure S2

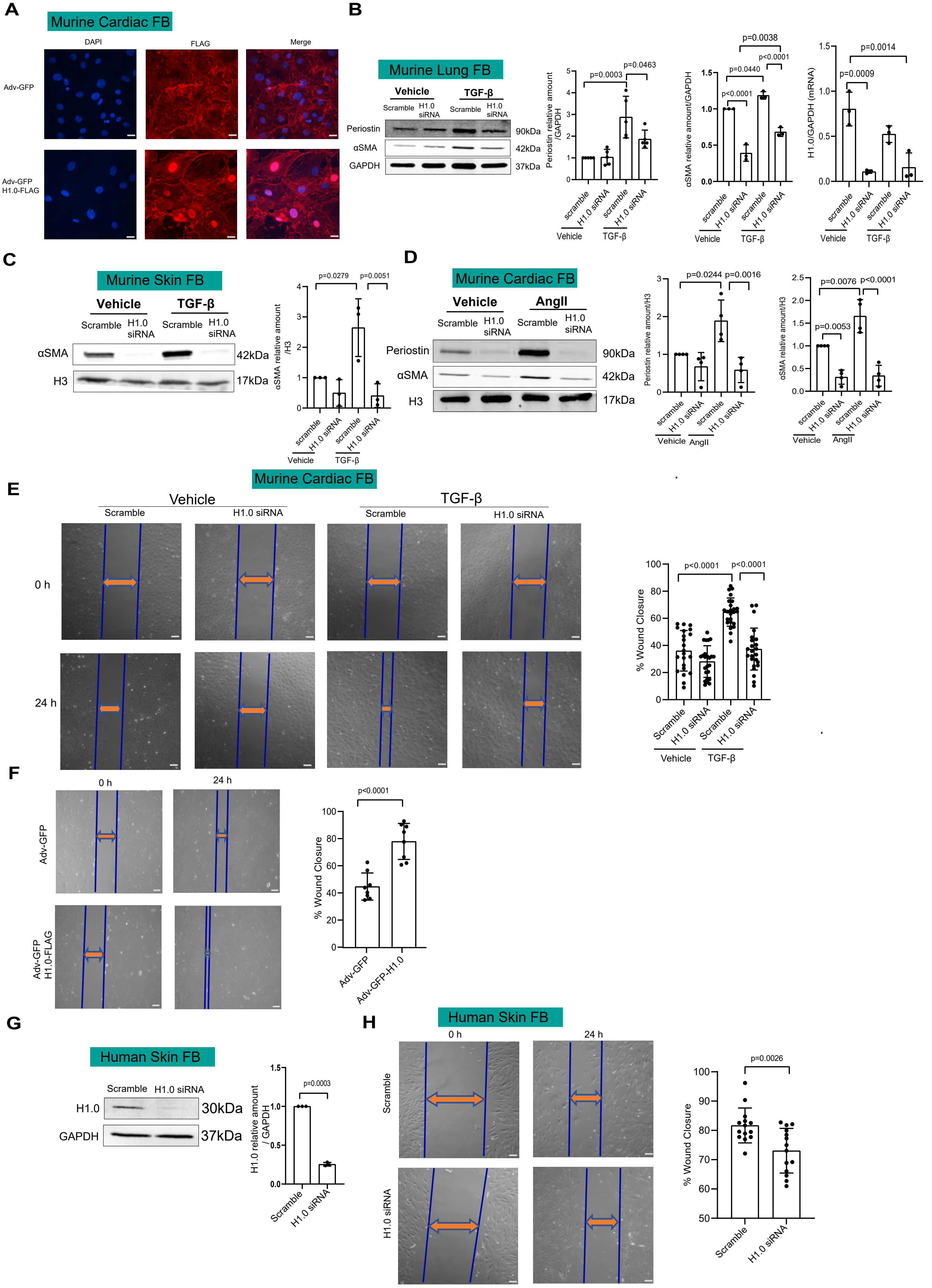
A
Murine Cardiac FB
DAPI
FLAG
Merge
Adv-GFP
Adv-GFP H1.0-FLAG
B
Murine Lung FB
Vehicle
TGF-β
Scramble
H1.0 siRNA
Periostin
αSMA
GAPDH
90kDa
42kDa
37kDa
Periostin relative amount /GAPDH
p=0.0003
p=0.0463
αSMA relative amount/GAPDH
p=0.0038
p=0.0440
p<0.0001
H1.0/GAPDH (mRNA)
p=0.0014
p=0.0009
scramble
H1.0 siRNA
C
Murine Skin FB
H3
17kDa
αSMA relative amount /H3
p=0.0279
p=0.0051
D
AngII
Periostin relative amount/H3
p=0.0244
p=0.0016
αSMA relative amount/H3
p=0.0076
p=0.0053
E
0 h
24 h
% Wound Closure
F
Adv-GFP
Adv-GFP H1.0-FLAG
Adv-GFP-H1.0
G
Human Skin FB
H1.0
30kDa
H1.0 relative amount / GAPDH
p=0.0003
H
p=0.0026

Figure S3

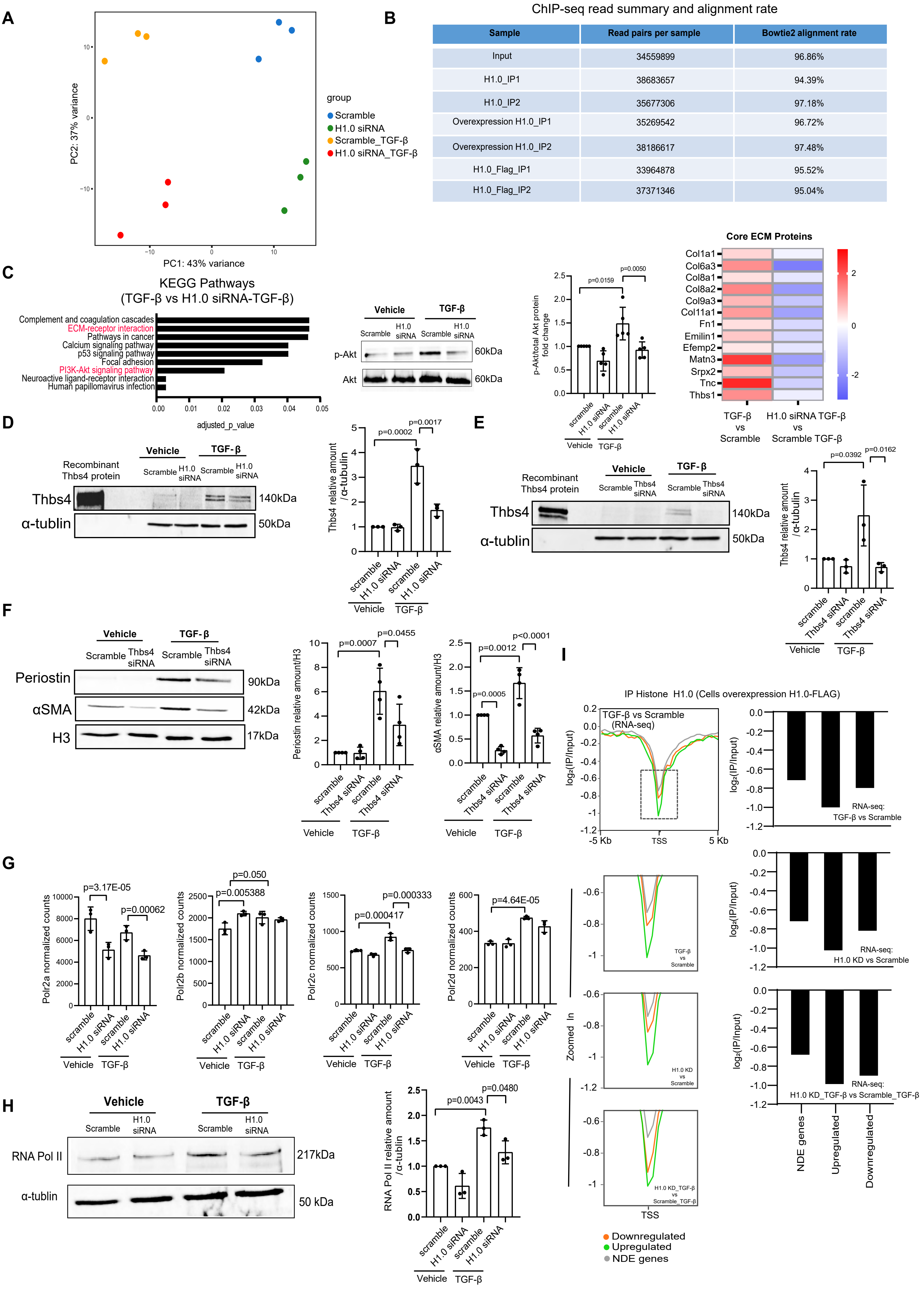


ChIP-seq read summary and alignment rate

| Sample | Read pairs per sample | Bowtie2 alignment rate |
|---|---|---|
| Input | 34559899 | 96.86% |
| H1.0_IP1 | 38683657 | 94.39% |
| H1.0_IP2 | 35677306 | 97.18% |
| Overexpression H1.0_IP1 | 35269542 | 96.72% |
| Overexpression H1.0_IP2 | 38186617 | 97.48% |
| H1.0_Flag_IP1 | 33964878 | 95.52% |
| H1.0_Flag_IP2 | 37371346 | 95.04% |

Figure S4

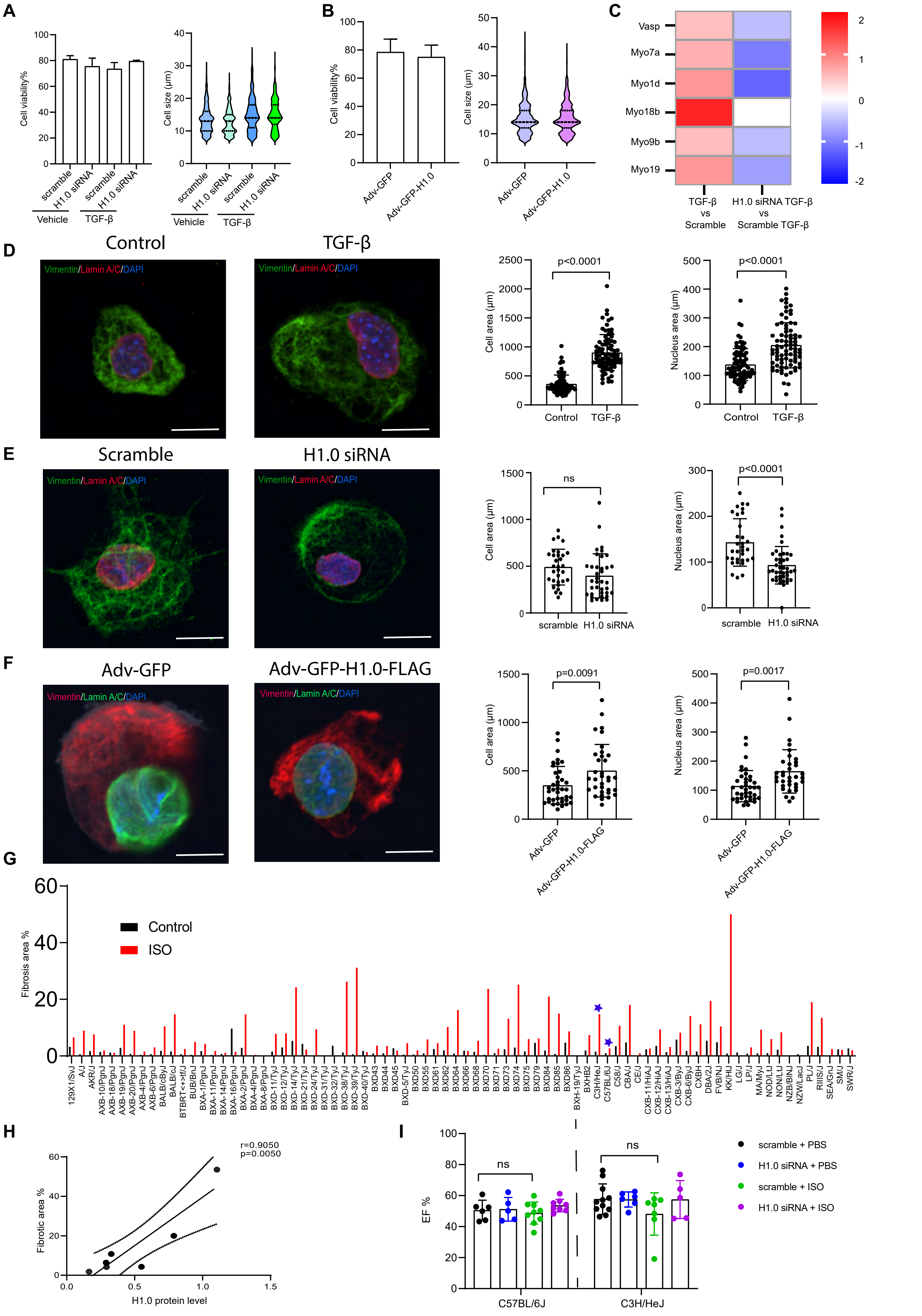
A
Cell viability%
scramble
H1.0 siRNA
Vehicle
TGF-β
Cell size (μm)
B
Adv-GFP
Adv-GFP-H1.0
C
Vasp
Myo7a
Myo1d
Myo18b
Myo9b
Myo19
TGF-β vs Scramble
H1.0 siRNA TGF-β vs Scramble TGF-β
D
Control
TGF-β
Vimentin/Lamin A/C/DAPI
Cell area (μm)
Nucleus area (μm)
p<0.0001
E
Scramble
H1.0 siRNA
ns
F
Adv-GFP
Adv-GFP-H1.0-FLAG
p=0.0091
p=0.0017
G
Fibrosis area %
Control
ISO
H
r=0.9050
p=0.0050
Fibrotic area %
H1.0 protein level
I
EF %
scramble + PBS
H1.0 siRNA + PBS
scramble + ISO
H1.0 siRNA + ISO
C57BL/6J
C3H/HeJ

Figure S5

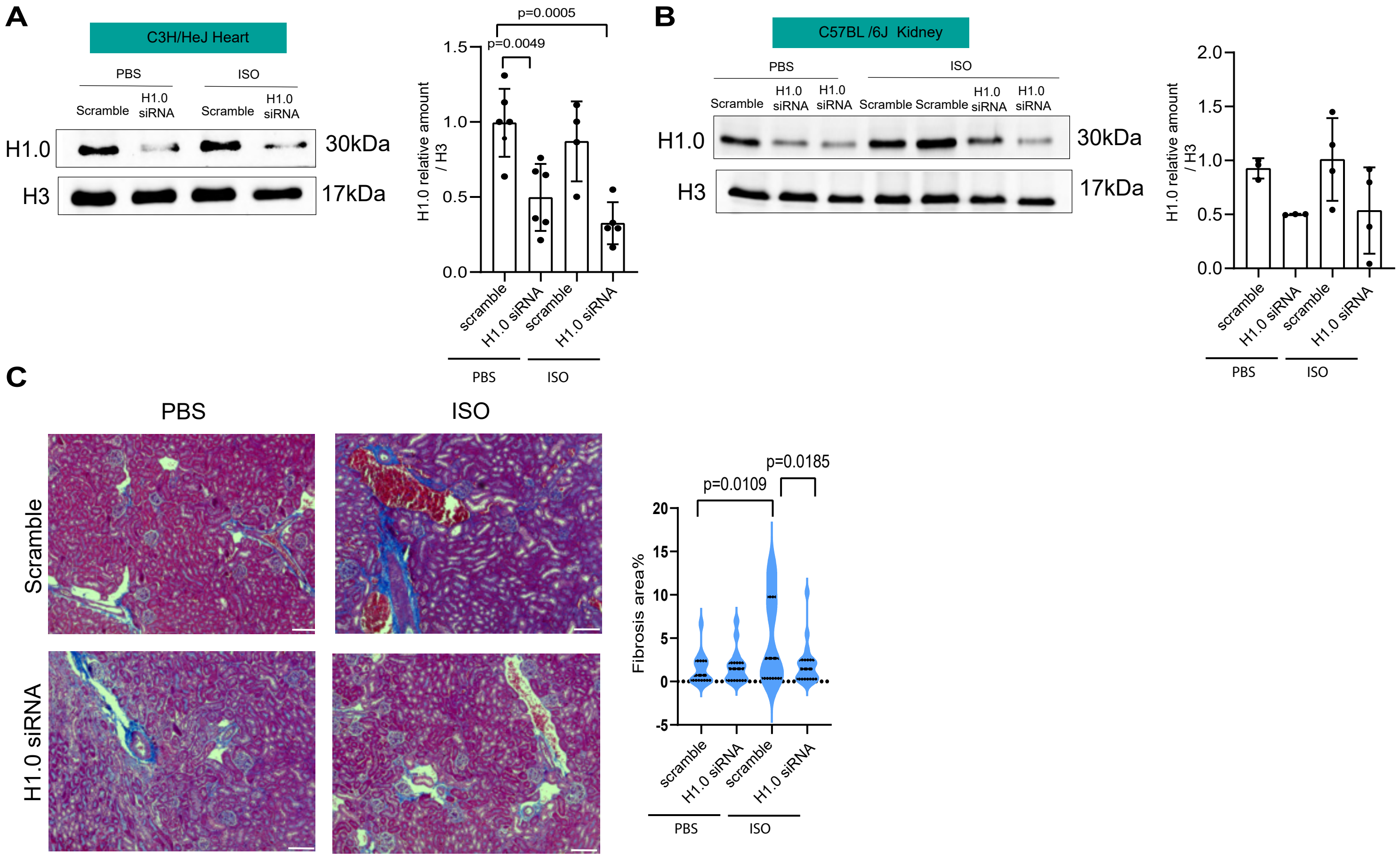
A
C3H/HeJ Heart
PBS
ISO
Scramble
H1.0 siRNA
Scramble
H1.0 siRNA
H1.0
H3
30kDa
17kDa
p=0.0005
p=0.0049
H1.0 relative amount / H3
scramble
H1.0 siRNA
scramble
H1.0 siRNA
PBS
ISO
B
C57BL /6J Kidney
PBS
ISO
Scramble
H1.0 siRNA
H1.0 siRNA
Scramble
Scramble
H1.0 siRNA
H1.0 siRNA
H1.0
H3
30kDa
17kDa
H1.0 relative amount / H3
scramble
H1.0 siRNA
scramble
H1.0 siRNA
PBS
ISO
C
PBS
ISO
Scramble
H1.0 siRNA
p=0.0109
p=0.0185
Fibrosis area%
scramble
H1.0 siRNA
scramble
H1.0 siRNA
PBS
ISO

Figure S6
